# Suppression of unwanted CRISPR/Cas9 editing by co-administration of catalytically inactivating truncated guide RNAs

**DOI:** 10.1101/597849

**Authors:** John C. Rose, Nicholas A. Popp, Christopher D. Richardson, Jason J. Stephany, Julie Mathieu, Cindy T. Wei, Jacob E. Corn, Dustin J. Maly, Douglas M. Fowler

## Abstract

CRISPR/Cas9 nucleases are powerful genome engineering tools, but unwanted cleavage at off-target and previously edited sites remains a major concern. Numerous strategies to reduce unwanted cleavage have been devised, but all are imperfect. Here, we report off-target sites can be shielded from the active Cas9•single guide RNA (sgRNA) complex through the co-administration of dead-RNAs (dRNAs), truncated guide RNAs that direct Cas9 binding but not cleavage. dRNAs can effectively suppress a wide-range of off-targets with minimal optimization while preserving on-target editing, and they can be multiplexed to suppress several off-targets simultaneously. dRNAs can be combined with high-specificity Cas9 variants, which often do not eliminate all unwanted editing. Moreover, dRNAs can prevent cleavage of homology-directed repair (HDR)-corrected sites, facilitating “scarless” editing by eliminating the need for blocking mutations. Thus, we enable precise genome editing by establishing a novel and flexible approach for suppressing unwanted editing of both off-targets and HDR-corrected sites.

## Introduction

The *S. pyogenes* Cas9 (SpCas9) nuclease is targeted to specific sites in the genome by a single guide RNA (sgRNA) containing a 20-nucleotide target recognition sequence. The target site must also contain an NGG protospacer adjacent motif (PAM)^1^. This multipartite target recognition system is imperfect, and most sgRNAs direct significant cleavage and subsequent unwanted editing at off-target sites whose sequence is similar to the target site^2–5^. Numerous approaches to reduce off-target editing have been devised, yet are hampered by various limitations^6–17^. For example, SpCas9 variants with improved specificity have been engineered^18–20^. While useful, these high-specificity variants often decrease on-target editing^21,22^ and in most cases do not eliminate all unwanted editing^20^. All high-specificity Cas9 variants appear to balance on- vs off-target activity via the same mechanism^20,23^ and, as a consequence, often fail to suppress editing at the same obstinate off-target sites^20,22^. Thus, new methods for off-target suppression are needed, particularly ones that preserve on-target editing, can be combined with high-specificity Cas9 variants, and require minimal expenditure of time, effort, and resources. To this end, we developed an orthogonal and general approach for suppressing off-targets that can be readily combined with existing methods, including high-specificity variants.

Our off-target suppression approach is based on the observation that sgRNAs with target recognition sequences 16 or fewer bases in length direct Cas9 binding to DNA target sites but do not promote cleavage^24–26^. Here, we show that Cas9 bound to dRNAs with perfect complementarity to off-target sites can dramatically improve editing specificity by shielding these sites from the active Cas9•sgRNA complex (**Fig. 1a**). To highlight the generality and ease of implementation of our method, which we call dRNA Off-Target Suppression (dOTS), we effectively suppress editing at 15 off-target sites, yielding up to a ∼40-fold increase in specificity, with minimal dRNA optimization. Furthermore, dOTS can be multiplexed to suppress several off-targets simultaneously and can be combined with other approaches for improving specificity. We also describe dRNA ReCutting Suppression (dReCS), wherein dRNAs prevent recutting of homology-directed repair (HDR)-corrected sites, eliminating the need for blocking mutations and facilitating “scarless” editing. Thus, we enable more precise genome editing by establishing a novel and flexible approach for suppressing unwanted editing of both off-target and HDR-corrected sites.

**Figure 1:**
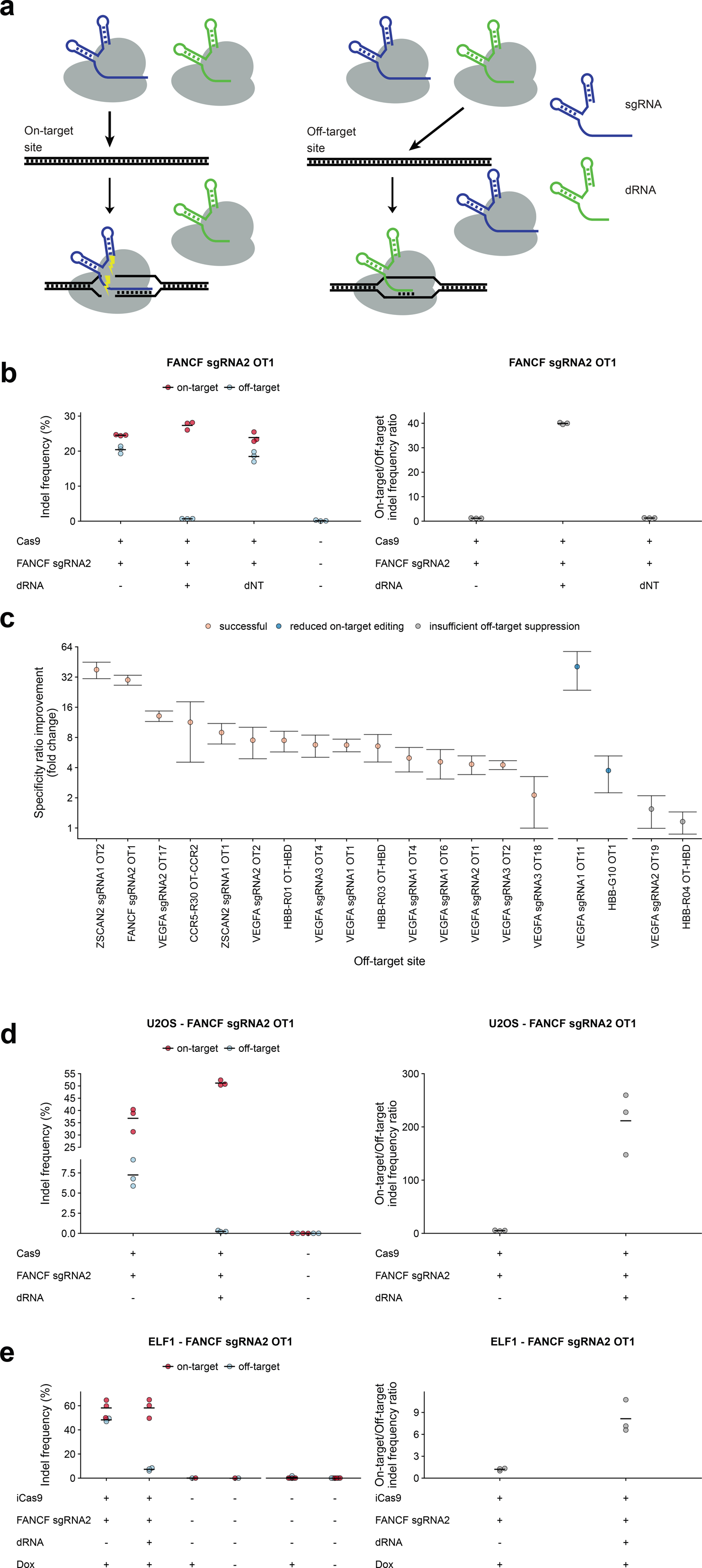
dRNA Mediated Off-Target Suppression (dOTS) effectively reduces off-target editing. **(a)** Schematic representation of dOTS. A dRNA (green) with perfect complementarity for an off-target site directs Cas9 binding but not cleavage, protecting the site. **(b)** Indel frequencies and specificity ratios (on-target/off-target indel frequency ratios) at the FANCF sgRNA2 on-target site and OT1 24 hours after transfection of HEK-293T cells with Cas9, sgRNA, and FANCF sgRNA2 OT1 dRNA1 or a non-targeting control dRNA (dNT) that does not target genomic DNA. For conditions without dRNA, an equivalent amount of pMAX-GFP was substituted. Means of n = 3 biological replicates depicted by solid lines. **(c)** Normalized specificity ratios, computed as the specificity ratio of the best dRNA condition (Supplementary Table 1) divided by the specificity ratio of the sgRNA only condition for 19 guide/off-target pairs tested in HEK-293T cells. Points depict the mean of n = 3 biological replicates, error bars show the standard error of the mean. OT = off-target. **(d)** Indel frequencies and specificity ratios at the FANCF sgRNA2 on-target site and OT1 24 hours after transfection in U2OS cells and **(e)** Elf1 embryonic stem cells. Control samples to the right of the x-axis break were performed separately. Means of n = 3 biological replicates depicted by solid lines.

## Results

### Dead RNA Off-Target Suppression (dOTS) increases on-target specificity

We first determined the feasibility of using dRNAs to suppress unwanted editing at off-target site 1 (OT1) of an sgRNA (sgRNA2) targeting the FANCF locus^18^. We co-transfected HEK-293T cells with a plasmid encoding SpCas9, along with equal amounts of plasmids encoding FANCF sgRNA2 and a GFP control, or FANCF sgRNA2 and one of four dRNAs with perfect complementarity to OT1 (**Fig. S1a**). Three of the four dRNAs significantly decreased off-target editing without appreciably impacting on-target editing, while co-transfection of a non-targeting control dRNA did not impact on-or off-target editing (**Fig. S1b**). In particular, dRNA1 decreased off-target editing from 20.44% (s.e.m. = 0.61%, n = 3) to 0.69% (s.e.m = 0.02%, n = 3), leading to a 30-fold increase in the on-target specificity ratio (**Fig. 1b**). Cas9•dRNA complexes are thought to lack cleavage activity, but a relatively small number of dRNAs have been evaluated so far^24,25^. Thus, we verified that dRNA1 did not direct any detectable Cas9 editing activity at either the on-or off-target sites (**Fig. S1c**).

To demonstrate the generality of dOTS, we evaluated 18 additional on-target/off-target pairs in HEK-293T cells. We found at least one dRNA for 15 of the 19 pairs we tested that increased the specificity ratio by at least two-fold (mean fold-change = 10.44) while decreasing on-target editing by no more than two-fold (mean fold-change = 0.93; **Fig. 1c, S2**). Across all on-target/off-target pairs, a median of six candidate dRNAs were screened, highlighting the ease of identifying effective dRNAs (**Fig. S2, Supplementary Table 1**). Non-targeting dRNAs did not impact editing (**Fig. S3**). Moreover, effective dRNAs did not induce indels at either on- or off-target sites, suggesting that few, if any, Cas9•dRNA complexes are active (**Supplementary Tables 2, 3**). dOTS was at least as effective in U2OS cells and the Elf1 naïve embryonic stem cell line as in HEK-293T cells (**Fig. 1d, e, S4**)^27^. Finally, we found that dRNA-mediated suppression of off-target editing was durable, with dRNAs effectively decreasing off-target editing for at least 72 hours post-transfection (**Fig. S5**).

An important application of Cas9 is editing genes containing pathogenic mutations^28,29^. For example, Cas9 has been used to cleave the *β*-globin locus (HBB), with the goal of knocking out sickle cell mutations^30,31^. However, the *δ*-globin locus (HBD) is a common off-target for sgRNAs targeting HBB, and cleavage of both on- and off-target sites can result in large chromosomal deletions at the globin locus^32^. In HEK-293T cells, dOTS decreased off-target editing at HBD from 1.08% (s.e.m. = 0.22%, n = 3) to 0.15% (s.e.m. = 0.03, n = 3; **Fig. S2d**). In Elf1 cells, dOTS decreased off-target editing at HBD from 20.72% (s.e.m. = 2.75, n = 3) to 1.20% (s.e.m. = 0.18, n = 3), increasing the specificity ratio from 1.33 to 13.72 (**Fig. S4b**). Thus, dOTS can control unwanted editing at clinically relevant loci.

We were unable to find effective dRNAs for four off-target sites. In two cases, dRNAs strongly reduced off-target editing but also decreased on-target editing by greater than two-fold (**Fig. 1c, S2b, i**). In two other cases, no dRNA we tested was effective in decreasing off-target editing (**Fig. 1c, S2e, m, n**). We suspect that these ineffective dRNAs are either unstable, form unfavorable secondary structures, or have insufficient affinity for the off-target site relative to their cognate sgRNAs. However, at most off-targets we identified one or more effective dRNAs that enhanced specificity without sacrificing on-target editing, making dOTS an effective approach for off-target suppression.

### Mechanism of off-target suppression by dRNAs

dOTS is based on our prediction that Cas9•dRNA complexes with perfect complementarity to an off-target site can directly outcompete active, imperfectly complementary Cas9•sgRNA complexes for binding. To test this Cas9 self-competition mechanism, we performed *in vitro* cleavage assays with linear DNA substrates and purified Cas9 ribonucleoprotein complexes (RNPs) containing either FANCF sgRNA2 or dRNA1. Incubation of a substrate containing the FANCF OT1 site with a mixture of the Cas9•dRNA1 and Cas9•sgRNA2 complexes led to a robust reduction in cleavage compared to administration of the Cas9•sgRNA2 complex alone (**Fig. 2a**). Consistent with our self-competition mechanism, preincubation of the substrate with the Cas9•sgRNA2 complex followed by addition of the Cas9•dRNA1 complex eliminated the reduction in cleavage (**Fig. S6a, b**). Thus, Cas9•dRNA complexes can directly shield off-target loci from Cas9•sgRNA cleavage.

**Figure 2:**
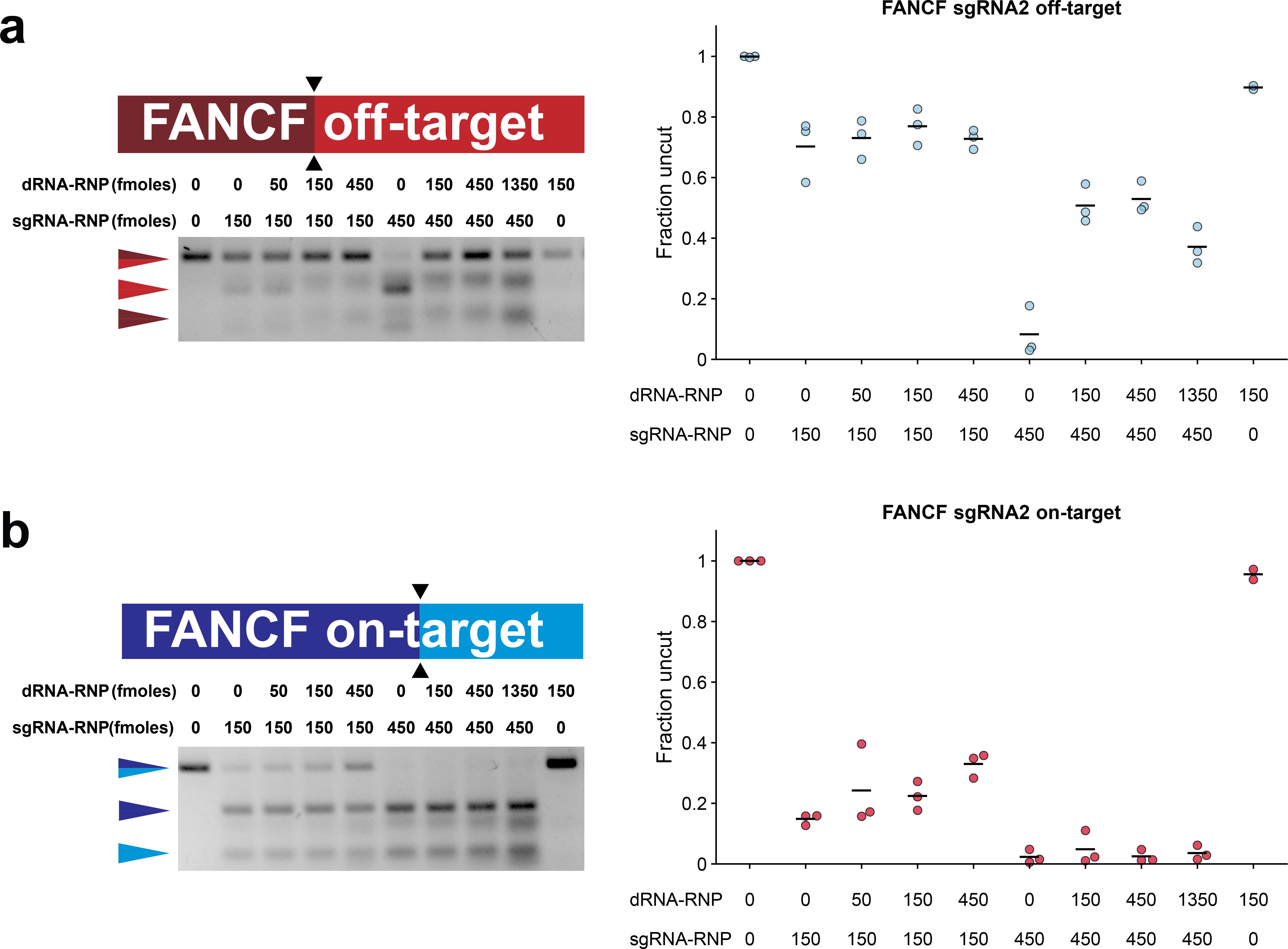
dRNAs suppress off-target editing by competing with sgRNAs for off-target sites. Representative gels of *in vitro* cleavage of PCR products containing either **(a)** FANCF sgRNA2 OT1 or **(b)** the FANCF sgRNA2 on-target site with either 150 or 450 fmoles of Cas9 FANCF sgRNA2 RNP in the presence of variable amounts of the Cas9 FANCF sgRNA2 OT1 dRNA1 complex. Fraction of uncut DNA determined by gel densitometry. Means of n = 3 replicates depicted by solid lines.

At low concentrations of Cas9•sgRNA2, Cas9•dRNA1 modestly reduced cleavage of the on-target FANCF substrate site *in vitro* (**Fig. 2b**), despite this dRNA not affecting on-target editing efficiency in cells (**Fig. 1b, d, e**). One possible explanation for this disparity is that, in cells, Cas9•dRNA1-mediated protection of the on-target locus decreases the rate of indel formation but editing reaches the same maximum as in cells without dRNA1 by the time of measurement. Another explanation is that cellular factors prevent Cas9•dRNA1, which should have modest affinity for the on-target site, from providing appreciable protection from cleavage by Cas9•sgRNA2. Thus, we measured rates of indel formation at FANCF sgRNA2 OT1 and the on-target site in cells using a chemically-inducible Cas9 (ciCas9) variant^6,33^. The activity of ciCas9 is repressed by an intramolecular autoinhibitory switch. Addition of a small molecule, A-1155463 (A115), disrupts autoinhibition and rapidly activates ciCas9, enabling precise studies of editing kinetics.

As expected, activation of ciCas9 with A115 led to the rapid appearance of indels at the FANCF sgRNA2 on- and off-target sites in the absence of dRNA1. Inclusion of a plasmid encoding dRNA1 effectively eliminated ciCas9-mediated editing at the off-target site but had no measurable impact on the kinetics of on-target editing (**Fig. 3a, S6c**). These results suggest that dRNAs with imperfect complementarity to an on-target site can bind to and protect that site in cell-free systems, but not in cells. The most likely explanation for this difference is that, in cells, DNA is subject to a variety of active processes that influence Cas9^34,35^. For example, the degree of complementarity between a guide and its target affects the ability of polymerases to displace dCas9 from DNA^36^, suggesting that polymerases may limit the ability of imperfectly complementary Cas9•dRNA complexes to shield on-target sites.

**Figure 3:**
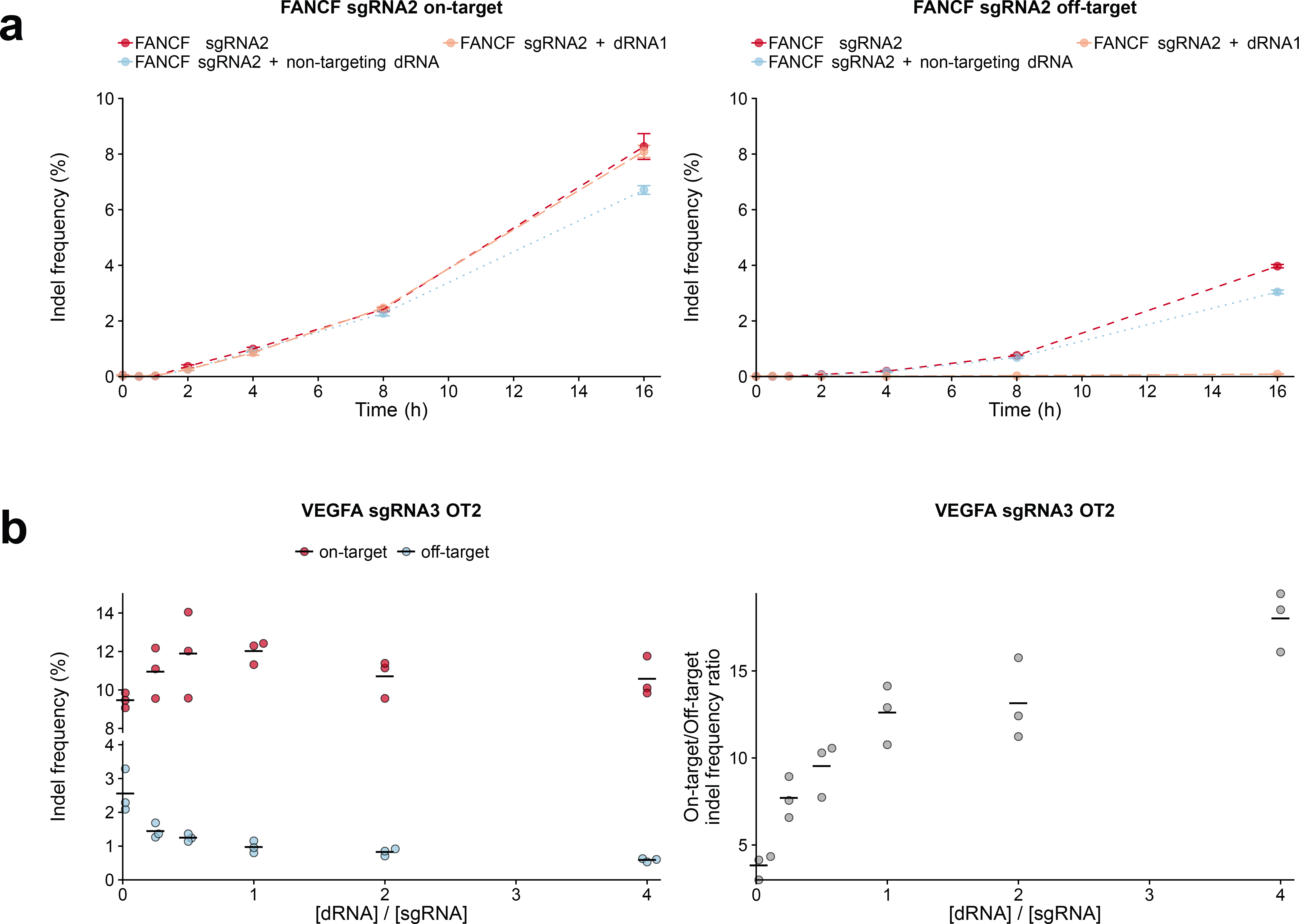
dRNAs affect off-target, but not on-target, editing kinetics and can be titrated to improve specificity. **(a)** Editing of FANCF sgRNA2 on-target and OT1 sites using chemically inducible Cas9 (ciCas9) from 0 to 16 hours after activation with A115. dNT is a 14-base control dRNA targeting a non-endogenous site. Points depict the mean of n = 3 biological replicates. Error bars show the standard error of the mean. **(b)** Indel frequencies and specificity ratios at VEGFA sgRNA3 on-target and OT2 sites in cells transfected with plasmids encoding Cas9 and varying ratios of VEGFA sgRNA3 and dRNA2. dRNA untreated cells were transfected with Cas9 and a 1:1 VEGFA sgRNA3:GFP plasmid ratio. Means of n = 3 biological replicates depicted by solid lines. OT = off-target.

Our proposed Cas9 self-competition mechanism predicts that the level of off-target shielding provided by moderately effective dRNAs can be improved by manipulating the ratio of Cas9•dRNA to Cas9•sgRNA in cells. While an initial 1:1 plasmid ratio was effective for all 15 successful dRNAs, increasing the amount of dRNA relative to sgRNA further decreased off-target editing and improved the specificity ratio at each of the four sgRNA/dRNA pairs we tested (**Fig. 3b, S7**). For one pair, higher dRNA:sgRNA ratios also decreased on-target editing. Thus, a trade-off between maintaining on-target editing and decreasing off-target editing exists for some sgRNA/dRNA pairs. Here, the dRNA/sgRNA ratio can be tuned based on whether preserving on-target editing or suppression of a particular off-target is desired.

### Combining dOTS with other approaches to improve Cas9 specificity

Other strategies to improve Cas9 specificity fail to completely suppress off-target editing and often reduce on-target efficacy. Thus, we wondered whether they could be enhanced with dOTS. One approach uses truncated sgRNAs (tru-sgRNAs) with 17-19 base target sequences to increase on-target specificity at some loci. For example, truncation of the VEGFA sgRNA3 target sequence (VEGFA tru-sgRNA3) decreases editing at some off-target sites, but editing at OT2 remains^11^. dOTS suppressed editing at this refractory off-target site without affecting on-target editing (**Fig. 4a**), demonstrating that it is compatible with tru-sgRNAs.

**Figure 4:**
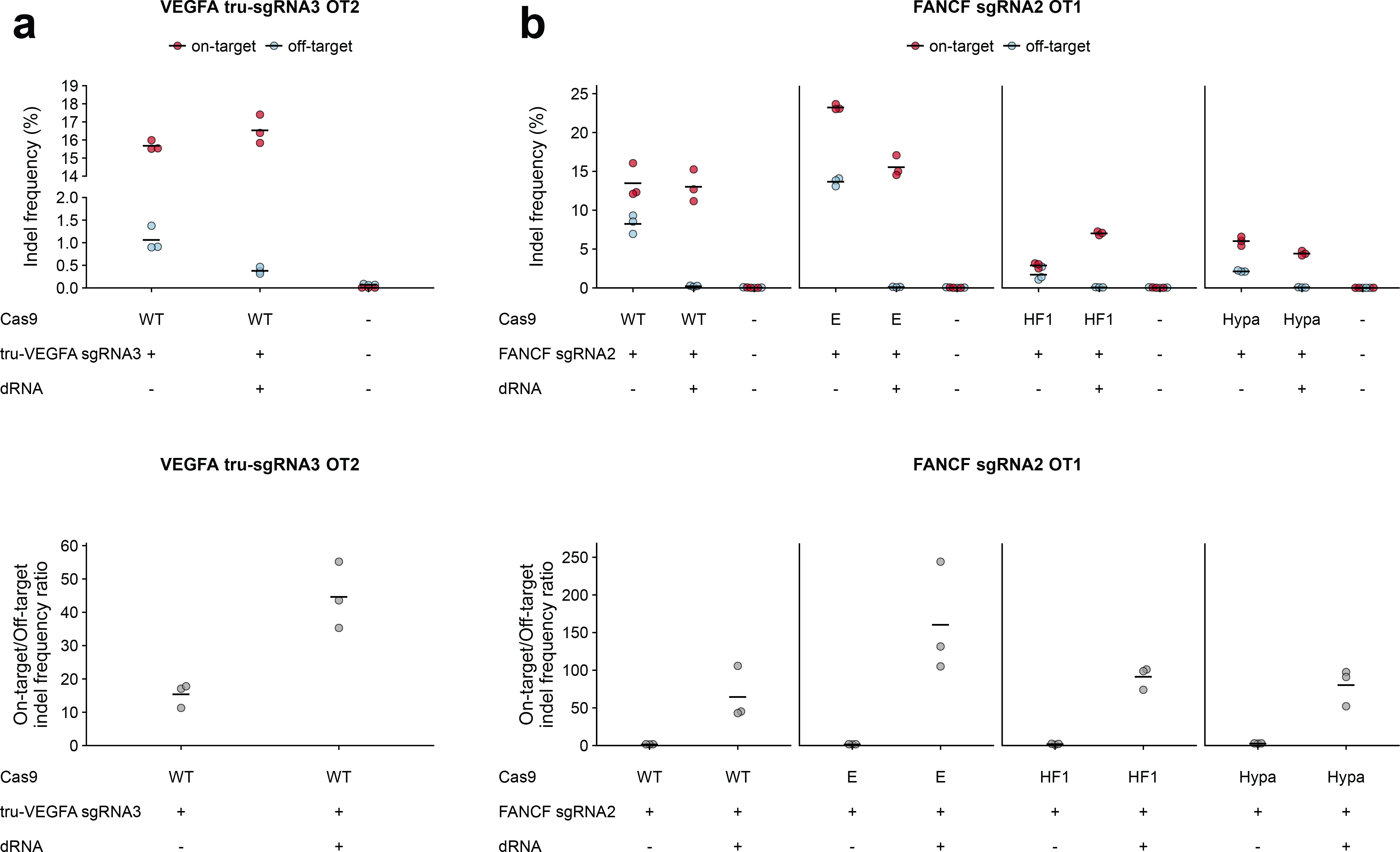
dRNAs can be combined with other approaches for improving Cas9 specificity. Indel frequencies and specificity ratios 24 hours after transfection with **(a)** plasmids encoding WT Cas9, a dRNA targeting VEGFA sgRNA3 OT2 (dRNA2) and a truncated guide VEGFA tru-sgRNA3 or **(b)** High-specificity variants of Cas9 and a dRNA targeting FANCF sgRNA2 OT1 (dRNA1). WT = wildtype Cas9, E = eSpCas9, HF1 = SpCas9-HF1, Hypa = HypaCas9. Means of n = 3 biological replicates depicted by solid lines. OT = off-target.

More recently, rational engineering of SpCas9 has produced high-specificity variants like eSpCas9(1.1), SpCas9-HF1, and HypaCas9^18–20^. While these variants generally improve on-target specificity, they do not suppress unwanted editing at all off-target sites for all sgRNAs. For example, a recent evaluation of these three high-specificity variants revealed off-target editing by all three variants for four of the six sgRNAs tested^20^. In another example, FANCF sgRNA2 OT1 is still edited at high frequencies by all three high-specificity variants (**Fig. 4b**)^18,20^. Co-transfection of FANCF sgRNA2 with an effective dRNA reduced off-target editing to levels indistinguishable from non-transfected controls for all high-specificity Cas9 variants (*P* > 0.05, one-sided t-test, n = 3), dramatically increasing specificity ratios (**Fig 4b**). dRNAs also effectively suppressed off-target editing by eSpCas9(1.1) and SpCas9-HF1 at a refractory VEGFA sgRNA3 off-target (**Fig. S8**). Thus, dOTS can be combined with many other methods for improving Cas9 specificity.

### dOTS can suppress off-targets at multiple sites simultaneously

Since many sgRNAs induce off-target editing at numerous sites^4,5,37^, we examined whether dOTS could be multiplexed. We selected three off-target sites for VEGFA sgRNA2 with individually effective dRNAs (**Fig. 1c, S2**). HEK-293T cells were transfected with VEGFA sgRNA2 and the dRNAs individually, in duplex, or in triplex. Even when all three dRNAs were combined, editing at each off-target site was suppressed by its cognate dRNA with only small losses in on-target editing (**Fig 5a, Fig. S9a**). Multiplex dOTS was also effective for two other sgRNAs (**Fig. S9b, c**), and could even suppress the off-targets of two distinct sgRNAs simultaneously (**Fig. S9d**).

**Figure 5:**
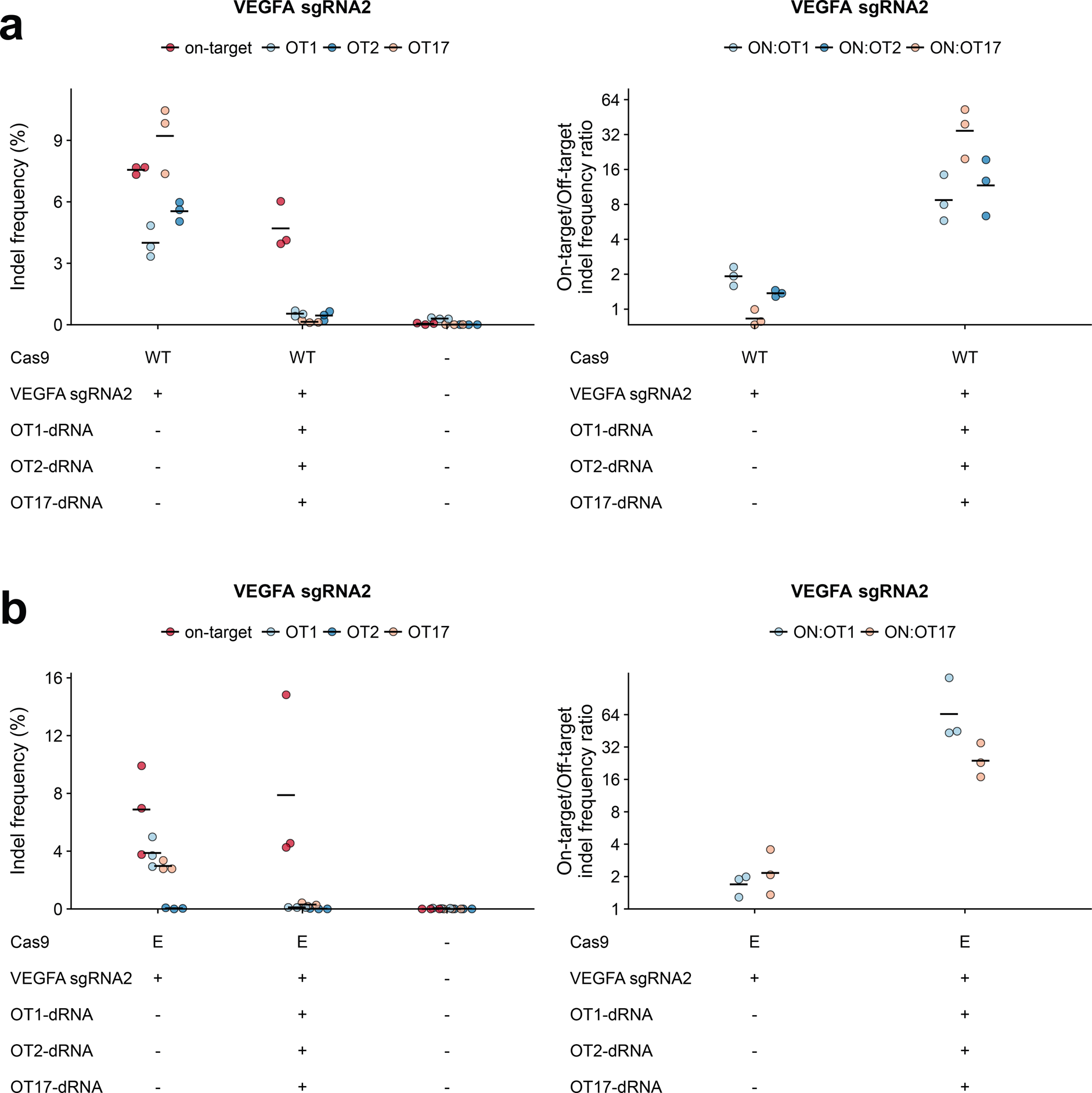
dRNAs can be multiplexed to suppress several off-targets simultaneously. Indel frequencies and specificity ratios 24 hours after transfection of plasmids encoding either **(a)** wild type (WT) or **(b)** eSpCas9 (E), VEGFA sgRNA2, and dRNAs targeting one of three VEGFA sgRNA2 off-targets (OT1 dRNA1, OT2 dRNA8, OT17 dRNA8). Means of n = 3 biological replicates depicted by solid lines. OT = off-target.

Like wild type Cas9, high-specificity Cas9 variants can cause editing at multiple off-target sites. For example, eSpCas9 reportedly drives appreciable editing with VEGFA sgRNA2 at three different off-target sites^20^. We observed off-target editing at two of these sites, and found that dRNAs could simultaneously decrease off-target editing at both sites without perturbing on-target editing (**Fig. 5b**). Furthermore, multiplexed dOTS suppressed editing driven by SpCas9-HF1 and HypaCas9 at these off-target sites (**Fig. S10**). Thus, in the context of both wild type and variant Cas9, dRNAs can be combined to suppress multiple off-targets simultaneously.

### dRNAs enable scarless HDR-mediated genome editing

When mutations introduced by HDR do not substantially disrupt the target sequence or PAM, as is generally the case for single nucleotide variants, Cas9 can continue to cleave the target site after repair. Continued cleavage introduces indels, substantially decreasing the frequency of loci containing the desired sequence. For example, quantification of editing outcomes at PSEN1 revealed that up to 95% of HDR-corrected templates showed secondary indels due to recutting^38^. If a protein-coding region is being edited, synonymous blocking mutations that disrupt the sgRNA target sequence, PAM, or both are generally included in the repair template. Unfortunately, synonymous blocking mutations may alter protein expression or interfere with mRNA splicing. Furthermore, predicting functionally neutral blocking mutations in non-coding regions is extremely challenging. Thus, “scarless” editing, the ability to efficiently introduce single nucleotide variants and other small changes into the genome without blocking mutations or unwanted indels, would be of tremendous utility.

We predicted that dRNAs directed at a desired, HDR-corrected sequence could shield repaired sites from recutting, an approach we call dRNA-mediated Re-Cutting Suppression (dReCS; **Fig. 6a**). We evaluated the ability of dRNAs to improve the HDR-mediated conversion of BFP to GFP through substitution of a single amino acid. Previously, several blocking mutations were used to prevent recutting, yet only a single nucleotide change is needed to alter the His in BFP (**<u>C</u>**AT) to the Tyr in GFP (**<u>T</u>**AT)^39^. We selected a previously used sgRNA in which the permissive site within the PAM (*i.e.* N in NGG) for the BFP sgRNA corresponds to the mutated nucleotide. Thus, this sgRNA possesses perfect complementarity to both the native and HDR-repaired locus, representing a worst-case scenario in which Cas9•sgRNA is expected to efficiently recut HDR-repaired sites. HEK-293T cells with stably integrated BFP were transfected with a single stranded oligodeoxynucleotide (ssODN) donor template containing the single nucleotide change, the sgRNA targeting BFP, and one of three dRNAs with perfect complementarity to the GFP but not BFP sequence. After four days, in the absence of dRNA, scarless HDR conversion to GFP was inefficient, with 1.94% of cells expressing GFP by flow cytometry. In the presence of the best dRNA, absolute HDR efficiency increased to 3.77% (**Fig. 6b, S11**), corresponding to an increase in the percentage of all edited sites exhibiting scarless HDR from 9.53% (s.e.m. = 0.40, n = 3) to 19.72% (s.e.m. = 0.52, n = 3; **Fig. 6c**). Thus, dReCS can promote scarless HDR even when the sgRNA has perfect complementarity for the HDR corrected sequence.

**Figure 6:**
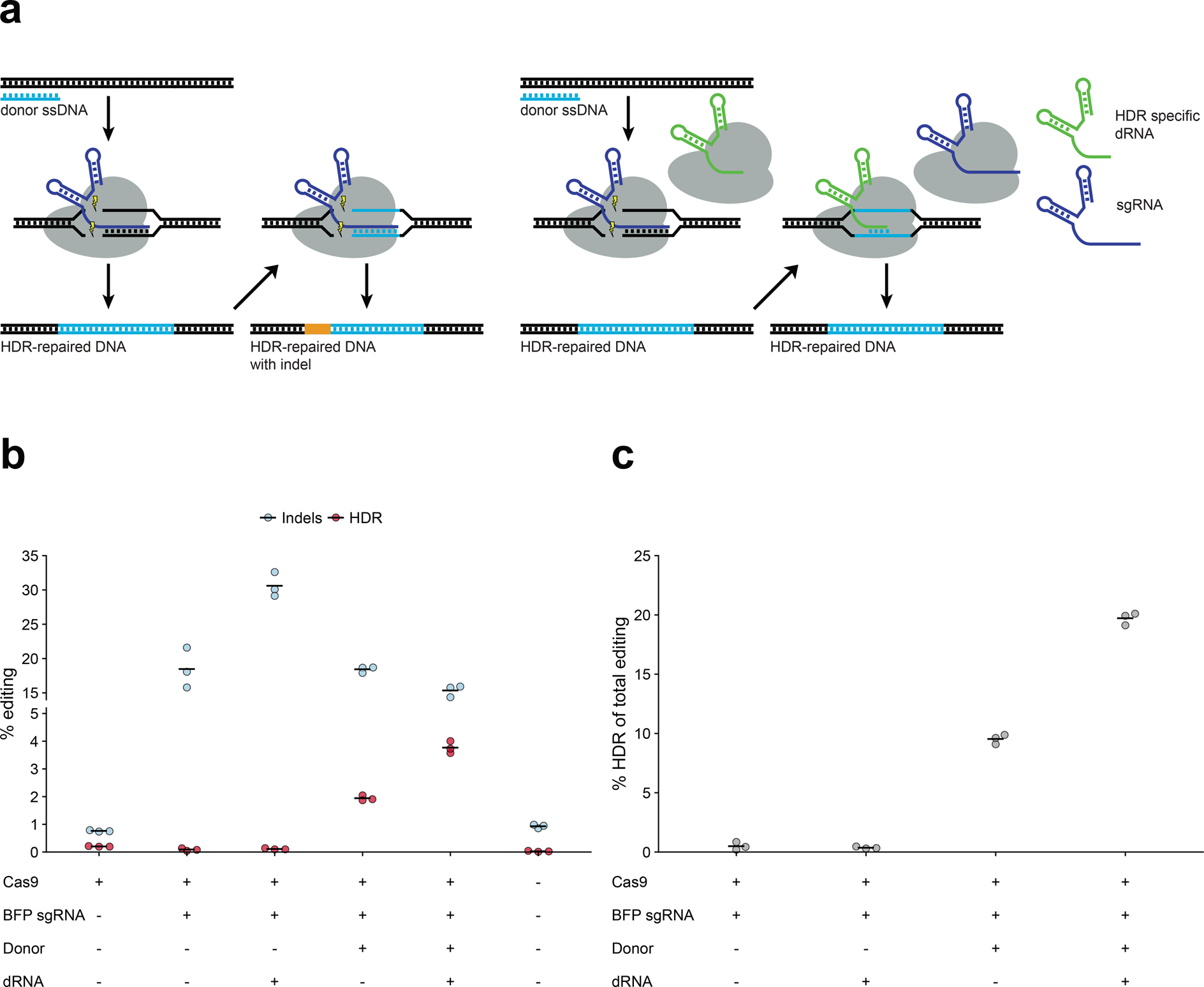
dRNA On-target Recutting Suppression (dReCS) facilitates scarless HDR. **(a)** Schematic depicting dReCS. dRNA (green) exhibiting perfect complementarity for the repaired site directs Cas9 binding but not cleavage, protecting the repaired site. **(b)** Indels and homology-directed repair (HDR) as assessed by flow cytometry, where indels lead to a loss of BFP signal, and HDR leads to a loss of BFP and gain of GFP signal. **(c)** HDR as a percentage of total Cas9 edits observed. Means of n = 3 biological replicates depicted by solid lines. dRNA = BFP sgRNA1 dRNA3 (see **Fig. S11**, **Supplementary Data Set 1**).

## Discussion

Here, we describe a general approach for the targeted suppression of unwanted Cas9-mediated editing that relies on co-administration of dRNAs with complementarity to the suppressed site. Our approach exploits the previously unappreciated phenomenon we refer to as Cas9 self-competition: the ability of different Cas9•guide RNA complexes to compete for a limited number of genomic target sites. We show that catalytically inactive Cas9, in this case Cas9 bound to a dRNA, can protect sites from undesired cleavage by active Cas9•sgRNA complexes. One application of this approach, dRNA mediated off-target suppression (dOTS), reduced editing at 15 distinct off-target sites, in some cases below the limit of detection by high-throughput sequencing. Another application, dRNA recutting suppression (dReCS), facilitated the scarless introduction of a single base change that did not impact the PAM or target sequence. dReCS circumvents the need for blocking mutations, making it particularly useful for single nucleotide variants and small indels in non-coding regions of the genome where synonymous blocking mutations are not an option. In both cases, effective dRNAs can generally be rapidly identified with minimal screening. Moreover, dRNAs are effective in a variety of different cell lines and they can be combined to protect multiple off-target sites simultaneously.

dOTS and dReCS offer many advantages, but they are not perfect. We could not find an effective dRNA for four of the 19 target/off-target pairs we tested. In some cases, additional dRNAs could be screened, but the sequence restrictions imposed by the SpCas9 NGG PAM mean that effective dRNAs may not always exist. One alternative is to improve poorly performing dRNAs by manipulating dRNA/sgRNA ratios. Another is to combine dRNAs with the recently described xCas9 variant, which has a more permissive PAM that increases the number of candidate dRNAs^40^. Another drawback is that some dRNAs decrease on-target editing, particularly when they are multiplexed to suppress several off-target sites simultaneously. We suspect that these losses in on-target editing likely arise due to dilution of the plasmids encoding the on-target sgRNA and/or Cas9, and could be reduced by transfecting ribonucleoprotein mixtures^41^ or using a multiplex guide expression scheme^42,43^. Finally, dRNAs could yield unwanted transcriptional off-target effects. However, transcriptional repression by Cas9 in the absence of a repressive domain is modest^44,45^, and such effects would be transient unless both Cas9 and the dRNA were integrated into the genome.

Other approaches for minimizing off-target editing are also imperfect, as they reduce on-target efficiency^6–9,21,22^, introduce new off-target sites^11,14,15^, limit the number of potential target sites^11,14–17^, or demand difficult Cas9 engineering^18–22,46,47^. Moreover, many of these approaches are laborious to implement in experimental models where Cas9 or a variant thereof has already been stably integrated into the genome^6–9,16–22,46,47^. Finally, these existing methods are generally incompatible with each other, meaning they cannot be used in concert to minimize limitations and improve performance. In contrast, dOTS and dReCS are comparatively easy to use, low-cost, and flexible. For example, dOTS could be used to address refractory off-targets of the popular engineered high-specificity Cas9 variants^18–22,46,47^. Here, we showed that dOTS could effectively suppress editing at four refractory off-target sites with three high-specificity Cas9 variants. Using dOTS to address these refractory off-targets is also far less laborious and time-intensive than further Cas9 engineering, as has been done previously^18,46^. Additionally, dReCS is simpler and less time-consuming than CORRECT^38^, a previous approach for scarless HDR editing that requires multiple rounds of HDR to introduce and subsequently remove blocking mutations. Because of their flexibility and technical simplicity, dOTS and dReCS could be readily integrated with existing protocols and experimental systems, enabling refinement of genome editing with minimal effort.

The flexibility of dOTS and dReCS means that they have applications beyond those we demonstrated. For instance, dOTS could facilitate allele-specific editing, even when the two alleles cannot be distinguished by a Cas9•sgRNA complex alone. Based on the principle of Cas9 self-competition, electroporation of Cas9•dRNA RNPs to quench editing by the active Cas9•sgRNA RNP should allow fine tuning of editing efficiencies. Similarly, dOTS could be employed to modulate the editing rates in CRISPR lineage tracing^48^. Finally, dOTS and dReCS are likely to be effective with other CRISPR enzymes, such as SaCas9 or Cpf1. Thus, dOTS and dReCS are easy-to-implement, effective and complementary methods for refining genome editing in both research and clinical applications.

## Supporting information

Supplementary Data Set 1

## Acknowledgements

This work was supported by the NIH (R01GM086858 (D.J.M), R01GM109110 (D.M.F), F30CA189793 (J.C.R)) and NSF (0954242 (D.J.M.)). D.M.F. is a CIFAR Azrieli Global Scholar.

The authors would like to thank D. Prunkard for assistance with flow cytometry experiments and analysis, as well as C. Murray for advice and assistance in the conception of this project.

## Author Contributions

J.C.R. conceived the study. J.C.R., J.E.C., D.M.F, and D.J.M designed the experiments. J.C.R., N.A.P., C.D.R., J.M., C.T.W. and J.J.S. performed the experiments. J.J.S. prepared samples for high-throughput sequencing. J.C.R., N.A.P., D.M.F and D.J.M wrote and edited the manuscript. All authors approved the final manuscript.

## Competing financial interests

The authors declare no competing financial interests.

## Methods

### Expression plasmids

All sgRNA and dRNA target sequences, except for VEGFA sgRNAs, were cloned into the gRNA_Cloning Vector according to the hCRISPR gRNA synthesis protocol (https://www.addgene.org/static/data/93/40/adf4a4fe-5e77-11e2-9c30-003048dd6500.pdf). gRNA_Cloning Vector was a gift from G. Church, Harvard (Addgene plasmid 41824). VEGFA site#1 (‘VEGFA sgRNA1’), VEGFA site#2 (‘VEGFA sgRNA2’) and VEGFA Site#3 (‘VEGFA sgRNA3’) were gifts from K. Joung, Massachusetts General Hospital (Addgene plasmids 47505, 47506 and 47507).

An N-terminal FLAG tag sequence was appended via Gibson Assembly Cloning (New England Biosciences) to a human codon optimized Cas9 (subcloned from hCas9, a gift from G. Church, Harvard; Addgene plasmid 41815) with a single C-terminal NLS expressed from a pcDNA3.3-TOPO vector. This was subsequently cloned into the pcDNA5/FRT/TO backbone (ThermoFisher). High-specificity variants of Cas9 — eSpCas9(1.1) (gift from F. Zheng, Broad Institiute; Addgene plasmid 71814) and VP12 (‘SpCas9-HF1’, gift from K. Joung, Massachusetts General Hospital; Addgene plasmid 72247) were subcloned into pcDNA5/FRT/TO backbone (ThermoFisher). HypaCas9 (‘BPK4410’) was a gift from J. Doudna and K. Joung, University of California, Berkeley and Massachusetts General Hospital (Addgene plasmid 101178).

The sequences of all plasmids, primers and other DNA constructs used in this work can be found in **Supplementary Data Set 1**.

### Cell culture

HEK-293T cells (293T/17, ATCC) were maintained in high-glucose DMEM supplemented with 10% fetal bovine serum (FBS, Life Technologies). U2OS cells (ATCC) were maintained in McCoy’s 5A (modified) medium supplemented with 10% FBS (Life Technologies). hESC Elf1 iCas9^27^ were plated into matrigel-coated 24-well plates and cultured in MEF-conditioned media supplemented with 2iL-I-F (GSK3i, MEKi, LIF, IGF, bFGF). All cell lines were regularly tested and confirmed free from mycoplasma contamination.

### Genome editing by Cas9

Unless otherwise specified, HEK-293T cells were plated in 24-well plates at 1.5 × 10^5^ cells/well. The day after plating, cells were transfected with Turbofectin 8.0 (Origene). For all dOTS experiments, 1.5 *μ*L of Turbofectin 8.0 and 500 ng of plasmid DNA were transfected. For dRNA screening experiments, the plasmid DNA mixture contained 250 ng Cas9 (eSpCas9, Cas9-HF1, or HypaCas9), 125 ng sgRNA, and 125 ng dRNA. For wells without dRNA, the 125 ng of pMAX-GFP was substituted for the dRNA plasmid as a transfection control. For multiplex dOTS experiments, the plasmid DNA mixture contained 250 ng Cas9, 125 ng sgRNA, and 125 ng each of 1-3 dRNAs. A pMAX-GFP plasmid was used to increase total DNA transfected per well to 750 ng. U2OS cells were plated in 12-well plates at 7.5 × 10^4^ cells/well. The next day they were transfected with 3 *μ*L of Turbofectin 8.0 and a total of 1 *μ*g plasmid DNA (500 ng Cas9, 250 ng sgRNA, and 250 ng dRNA or pMAX-GFP plasmid). For titration experiments with all sgRNAs except VEGFA sgRNA3, HEK-293T cells were transfected with 1.5 *μ*L of Turbofectin 8.0 and 500 ng of plasmid DNA. This DNA mixture contained 250 ng Cas9. The remaining 250 ng of DNA was divided between sgRNA and dRNA at varying ratios such that the total DNA was kept constant across experiments (1:1, 125 ng each sgRNA and dRNA; 1:2, 83.3 ng sgRNA and 166.7 ng dRNA; 1:4, 50 ng sgRNA and 200 ng dRNA; 2:1, 166.7 ng sgRNA and 83.3 ng dRNA; and 4:1, 200 ng sgRNA and 50 ng dRNA). For wells without dRNA, 125 ng of pMAX-GFP plasmid was substituted for the dRNA plasmid as a transfection control. For titration experiments with VEGFA sgRNA3, HEK-293T cells were transfected as above, but the DNA mixture contained 166.5 ng Cas9, and the various sgRNA:dRNA ratios were as follows (1:1, 166.5 ng each sgRNA and dRNA; 1:2, 111 ng sgRNA and 222 dRNA; 1:4, 66.6 ng sgRNA and 266.4 ng dRNA; 2:1, 222 ng sgRNA and 111 ng dRNA; 4:1, 266.4 ng sgRNA and 66.4 ng dRNA). For wells without dRNA, 166.5 ng of pMAX-GFP plasmid was substituted for the dRNA plasmid as a transfection control.

To harvest HEK-293T and U2OS cells for dOTS experiments, 24 hours after transfection each well of a 24-well plate was resuspended by thorough pipetting with 400 *μ*L ice-cold DPBS. Resuspended cells were then spun at 1,500 × *g* for 10 min at 4°C. DPBS was then aspirated and cell pellets were stored at −80°C until genomic DNA isolation. For extended timepoint experiments, the same protocol was followed, except cells were passaged into a new 24 well plate after 24 hours after transfection and then subsequently harvested 48 hours after passaging.

Two days prior to plating, hESC Elf1 iCas9 cells were treated with 2 *μ*g/ml doxycycline to induce Cas9 expression. At day 0, 2.5 × 10^4^ cells were plated into each well of a 24-well plate with addition of fresh doxycycline (2 *μ*g/ml) and 10 *μ*M Rock inhibitor to promote cell survival. After 24 hours, cells were transfected with 3 *μ*L of Genejuice (EMD Millipore) and 1 *μ*g plasmid DNA. This plasmid DNA mixture contained 500 ng sgRNA and 500 ng dRNA. For wells without dRNA, 500 ng of pMAX-GFP was substituted as a transfection control.

For Elf1 cells, 48 hours after transfection, each well of a 24-well plate was rinsed once with 0.5 mL DPBS and incubated for 5 min with trypsin to detach cells. 5 mL hESC media was added and the cells were spun down at 290 × *g* for 3 min. The pellet was then washed with 1 mL DPBS, spun again at 290 × *g* for 3 min then flash frozen in liquid nitrogen and stored at −80°C until genomic DNA isolation.

### dRNA recutting suppression (dReCS)

For dReCS experiments, a HEK-293T cell line with a genomically encoded BFP/GFP reporter was used^39^. The BFP/GFP reporter HEK-293T cell line contains a BFP that is converted to GFP via HDR-mediated substitution of a single amino acid (His in BFP (**<u>C</u>**AT) to Tyr in GFP (**<u>T</u>**AT)). BFP/GFP reporter cells were plated at 3.0 × 10^5^ cells/well in 12-well plates. 18 hours after plating, cells were transfected with 3 *μ*L of Turbofectin 8.0 (Origene) and 1,000 ng of total DNA. The total DNA mixture contained 272.7 ng of plasmid encoding Cas9, 54.5 ng sgRNA plasmid, 218 ng dRNA plasmid, and 454.5 ng symmetric or asymmetric single stranded donor DNA (**Supplementary Data Set 1**)^39^. For controls missing one or more of these DNA elements, the appropriate amount of DNA was replaced with a pKan-mCherry plasmid. Cells were maintained with standard passaging procedures for 4 days post-transfection until analysis by flow cytometry.

After 4 days, cells were washed with 2 mL DPBS, trypsinized with 0.5 mL 0.25% trypsin-EDTA (Life Technologies) for 2-4 minutes, and quenched with DMEM supplemented with 10% FBS. Cells were then spun down at 290 × *g* for 4 min, aspirated, and resuspended in DPBS supplemented with 1% FBS. Cells were run through a 35 *μ*m filter and analyzed by flow cytometry on an LSR-II flow cytometer. After gating for live cells (FSC-A vs SSC-A) and single cells (FSC-A x SSC-W), cells were analyzed for their BFP and GFP fluorescence. Gates for BFP and GFP positivity were determined by comparison to an untransfected BFP cell line. BFP+ GFP− cells were considered wildtype (WT). BFP− GFP− cells were considered to have undergone NHEJ but not HDR, as indels in this region of BFP lead to loss of fluorescence. Any cell that was GFP+ (regardless of residual BFP fluorescence) was considered to have undergone successful HDR. Percentages for each result (WT, HDR, NHEJ) were calculated as a fraction of the total cells that passed singlet gating. Percent HDR of total editing was determined as the fraction of cells with successful HDR divided by the total number of cells that underwent either HDR or NHEJ.

### *In vitro* Cas9 RNP nuclease assays

Cas9-2NLS in a pMJ915 vector (Addgene plasmid 69090) was expressed in *E. coli* and purified by a combination of affinity, ion exchange, and size exclusion chromatography as previously described^49^, except the final purified protein was eluted into a buffer containing 20 mM HEPES KOH pH 7.5, 5% glycerol, 150 mM KCl, 1 mM DTT at a final concentration of 40 *μ*M of Cas9-2NLS. FANCF sgRNA2 and FANCF dRNA1 were generated by HiScribe (NEB E2050S) T7 in vitro transcription using PCR-generated DNA as a template^49^, (dx.doi.org/10.17504/protocols.io.dm749m). Complete sequences for all sgRNA templates can be found in **Supplementary Data Set 1**.

A 463 basepair fragment containing the on-target cut site of FANCF sgRNA2 (FANCF target site) was PCR amplified from a custom FANCF sgRNA2 target site substrate gBlock (IDT) using primers oCR1711 and oCR1712. A 329 basepair fragment containing the cut site for off-target 1 of FANCF sgRNA2 (FANCF off-target) was PCR amplified from a custom FANCF sgRNA2 off-target substrate gBlock (IDT) using oCR1713 and oCR1714 (**Supplementary Data Set 1**). Prior to nuclease experiments, sgRNA and dRNA RNP complexes were generated by incubating purified Cas9-2NLS and FANCF sgRNA2 or dRNA1 in equimolar amounts for 10 minutes. For dRNA-RNP titration experiments, 150 or 450 fmoles of FANCF-sgRNA2-RNP complex and 0, 50, 150, or 450 fmoles of dRNA-RNP Cas9-sgRNA complex were co-added to 150 fmoles of FANCF target site or FANCF off-target substrate DNA. Reaction mixtures were incubated at 37 °C for 20 minutes in 20mM Tris, 100mM KCl, 5 mM MgCl_2_, 1 mM DTT, 0.01% Tween, 50 *μ*g/mL Heparin. Reactions were stopped by the addition of 1:4 volume of STOP solution (8mM Tris, 0.025% BPB, 0.025% XC, 50% Glycerol, 110mM EDTA, 1% SDS, 3mg/mL Proteinase K), followed by incubation at 55 °C for five minutes to liberate cut DNA fragments. Each digestion reaction was run on a 2% TAE agarose gel, post-stained with Ethidium Bromide, and resolved on a Gel-Doc (BioRad).

For pre-incubation experiments, FANCF sgRNA2 or dRNA1 RNP complexes were generated as described above. 450 fmoles of a single RNP complex was added to 150 fmoles of FANCF target site or FANCF off-target substrate DNA and incubated at 37 °C for 10 minutes. After 10 minutes, 450 fmoles of the other Cas9-RNP complex was added and allowed to incubate at 37 °C for an additional 10 minutes. Reactions were quenched, incubated, and run on a gel in an identical manner to the above experiments.

Gel densitometry analysis was performed in ImageJ. For each lane, background density was subtracted from the quantification of each band. The density of the uncut band was then divided by the total intensity of all bands in the lane to determine the uncut DNA fraction.

### Genomic editing by ciCas9

HEK-293T cells were treated according to previous methods^6^. Briefly, HEK-293T cells were plated in 12 well plates at 3.0 × 10^5^ cells/well. The day after plating, cells were transfected with 1.5 *μ*L Turbofectin 8.0 and 500 ng of plasmid DNA. The plasmid DNA mixture contained 250 ng Cas9, 125 ng FANCF sgRNA2 sgRNA, and 125 ng dRNA. For wells without dRNA, the 125 ng of dRNA plasmid were replaced by pMAX-GFP as a transfection control.

24 hours after transfection, cells were treated with with 10 *μ*M A115 dissolved in DMSO to induce ciCas9 activity. 24 hours after treatment with A115, cells were harvested after washing with 600 *μ*L DPBS to remove excess A115 and then resuspending cells in 600 *μ*L ice-cold DPBS. Resuspended cells were then spun at 1,500 × *g* for 10 min at 4°C. DPBS was aspirated and the cell pellets were stored at −80°C until genomic DNA isolation.

### Insertion and deletion detection by high throughput sequencing

Genomic DNA isolation, sequencing, and analysis were performed as previously described^6^. Briefly, genomic DNA was isolated using the DNEasy Blood and Tissue Kit (Qiagen) according to the manufacturer’s instructions except that the proteinase K digestion was conducted for 1 hr at 56°C. 15 cycles of primary PCR to amplify the region of interest was performed using 2 *μ*L of DNeasy eluate (∼100–300 ng template) in a 5 *μ*L Kapa HiFi HotStart polymerase reaction (Kapa Biosystems; for primers see **Supplementary Data Set 1**). The PCR reaction was diluted with 35 *μ*L DNAse-free water (Ambion). Illumina adapters and indexing sequences were added via 20 cycles of secondary PCR with 3 *μ*L of diluted primary PCR product in a 10 *μ*L Kapa Robust HotStart polymerase reaction (New England Biosciences; for primers see Supplementary Data Set 1). The final amplicons were run on a TBE-agarose gel (1.5%); and the product band was excised and extracted using the Freeze and Squeeze Kit according to the manufacturer’s instructions (Bio-Rad). Gel-purified amplicons were quantified using Qbit dsDNA HS Assay kit (Invitrogen). Then, up to 1200 indexed amplicons were pooled, quantified by Kapa Library Quantification (Kapa Biosysytems) and sequenced on a NextSeq (NextSeq 150/300 Mid V2 kit, Illumina, for primers see **Supplementary Data Set 1**).

Indels were quantified as previously described^6^. Briefly, after demultiplexing of reads (bcl2fastq/2.18, Illumina), indels were quantified with a custom Python script that is freely available upon request. 8-mer sequences were identified in the reference sequence located 20 bp upstream and downstream of the target sequence. Sequence distal to these 8-mers was trimmed. Reads lacking these 8-mers were discarded. For the VEGFA sgRNA3 OT2 locus, the process was the same, except 20-mer sequences located 10 bp upstream and downstream of the target sequence were used. For the VEGFA sgRNA3 OT4 locus, 8-mer sequences located 10 bp upstream and downstream of the target sequence were used. The trimmed reads were then evaluated for indels using the Python difflib package. Indels were defined as trimmed reads which differed in length from the trimmed reference and for which an insertion or deletion operation spanning or within 1 bp of the predicted Cas9 cleavage site was present. For dRNA only experiments, indels were quantified using both the sgRNA and dRNA predicted cut sites. Specificity ratios were calculated by dividing the indel percentage at the on-target locus by the indel percentage at the off-target locus for each sgRNA. For quantification of off-target editing for one of the VEGFA tru-sgRNA3 plus dRNA replicates (**Fig. 4a**), reads were acquired from multiple sequencing runs.

## Statistical Analysis

Statistical analysis of indel frequency and specificity ratios were performed using a one-sided two sample Student’s *t*-test.

## Data Availability Statement

Raw sequencing data will be made available upon publication through the NCBI GEO repository.

## References

1. Jiang, F. & Doudna, J. A. CRISPR-Cas9 Structures and Mechanisms. Annu Rev Biophys 46, 505–529 (2017).

2. Hsu, P. D. et al. DNA targeting specificity of RNA-guided Cas9 nucleases. Nat. Biotechnol. 31, 827–832 (2013).

3. Pattanayak, V. et al. High-throughput profiling of off-target DNA cleavage reveals RNA-programmed Cas9 nuclease specificity. Nat. Biotechnol. 31, 839–843 (2013).

4. Cameron, P. et al. Mapping the genomic landscape of CRISPR-Cas9 cleavage. Nat. Meth. 14, 600–606 (2017).

5. Tsai, S. Q. et al. GUIDE-seq enables genome-wide profiling of off-target cleavage by CRISPR-Cas nucleases. Nat. Biotechnol. 33, 187–197 (2015).

6. Rose, J. C. et al. Rapidly inducible Cas9 and DSB-ddPCR to probe editing kinetics. Nat. Meth. 14, 891–896 (2017).

7. Davis, K. M., Pattanayak, V., Thompson, D. B., Zuris, J. A. & Liu, D. R. Small molecule-triggered Cas9 protein with improved genome-editing specificity. Nat. Chem. Biol. 11, 316–318 (2015).

8. Zetsche, B., Volz, S. E. & Zhang, F. A split-Cas9 architecture for inducible genome editing and transcription modulation. Nat. Biotechnol. 33, 139–142 (2015).

9. Maji, B. et al. Multidimensional chemical control of CRISPR-Cas9. Nat. Chem. Biol. 13, 9–11 (2017).

10. Tycko, J., Myer, V. E. & Hsu, P. D. Methods for Optimizing CRISPR-Cas9 Genome Editing Specificity. Mol. Cell 63, 355–370 (2016).

11. Fu, Y., Sander, J. D., Reyon, D., Cascio, V. M. & Joung, J. K. Improving CRISPR-Cas nuclease specificity using truncated guide RNAs. Nat. Biotechnol. 32, 279–284 (2014).

12. Yin, H. et al. Partial DNA-guided Cas9 enables genome editing with reduced off-target activity. Nat. Chem. Biol. 14, 311–316 (2018).

13. Ryan, D. E. et al. Improving CRISPR-Cas specificity with chemical modifications in single-guide RNAs. Nucleic Acids Res. 46, 792–803 (2018).

14. Ran, F. A. et al. Double nicking by RNA-guided CRISPR Cas9 for enhanced genome editing specificity. Cell 154, 1380–1389 (2013).

15. Mali, P. et al. CAS9 transcriptional activators for target specificity screening and paired nickases for cooperative genome engineering. Nat. Biotechnol. 31, 833–838 (2013).

16. Guilinger, J. P., Thompson, D. B. & Liu, D. R. Fusion of catalytically inactive Cas9 to FokI nuclease improves the specificity of genome modification. Nat. Biotechnol. 32, 577–582 (2014).

17. Tsai, S. Q. et al. Dimeric CRISPR RNA-guided FokI nucleases for highly specific genome editing. Nat. Biotechnol. 32, 569–576 (2014).

18. Kleinstiver, B. P. et al. High-fidelity CRISPR-Cas9 nucleases with no detectable genome-wide off-target effects. Nature 529, 490–495 (2016).

19. Slaymaker, I. M. et al. Rationally engineered Cas9 nucleases with improved specificity. Science 351, 84–88 (2016).

20. Chen, J. S. et al. Enhanced proofreading governs CRISPR-Cas9 targeting accuracy. Nature 550, 407–410 (2017).

21. Vakulskas, C. A. et al. A high-fidelity Cas9 mutant delivered as a ribonucleoprotein complex enables efficient gene editing in human hematopoietic stem and progenitor cells. Nature Medicine 24, 1216–1224 (2018).

22. Lee, J. K. et al. Directed evolution of CRISPR-Cas9 to increase its specificity. Nat. Commun. 9, 3048 (2018).

23. Singh, D. et al. Mechanisms of improved specificity of engineered Cas9s revealed by single-molecule FRET analysis. Nat Struct Mol Biol 25, 347–354 (2018).

24. Kiani, S. et al. Cas9 gRNA engineering for genome editing, activation and repression. Nat. Meth. 12, 1051–1054 (2015).

25. Dahlman, J. E. et al. Orthogonal gene knockout and activation with a catalytically active Cas9 nuclease. Nat. Biotechnol. 33, 1159–1161 (2015).

26. Ye, L. et al. Programmable DNA repair with CRISPRa/i enhanced homology-directed repair efficiency with a single Cas9. Cell Discovery 2018 4:1 4, 46 (2018).

27. Ferreccio, A. et al. Inducible CRISPR genome editing platform in naive human embryonic stem cells reveals JARID2 function in self-renewal. cc 17, 535–549 (2018).

28. Ma, H. et al. Correction of a pathogenic gene mutation in human embryos. Nature 548, 413–419 (2017).

29. De Ravin, S. S. et al. CRISPR-Cas9 gene repair of hematopoietic stem cells from patients with X-linked chronic granulomatous disease. Sci. Transl. Med. 9, eaah3480 (2017).

30. Dever, D. P. et al. CRISPR/Cas9 β-globin gene targeting in human haematopoietic stem cells. Nature 539, 384–389 (2016).

31. DeWitt, M. A. et al. Selection-free genome editing of the sickle mutation in human adult hematopoietic stem/progenitor cells. Sci. Transl. Med. 8, 360ra134–360ra134 (2016).

32. Cradick, T. J., Fine, E. J., Antico, C. J. & Bao, G. CRISPR/Cas9 systems targeting β-globin and CCR5 genes have substantial off-target activity. Nucleic Acids Res. 41, 9584–9592 (2013).

33. Rose, J. C., Stephany, J. J., Wei, C. T., Fowler, D. M. & Maly, D. J. Rheostatic Control of Cas9-Mediated DNA Double Strand Break (DSB) Generation and Genome Editing. ACS Chem. Biol. 13, 438–442 (2018).

34. Clarke, R. et al. Enhanced Bacterial Immunity and Mammalian Genome Editing via RNA-Polymerase-Mediated Dislodging of Cas9 from Double-Strand DNA Breaks. Mol. Cell 71, 42–55.e8 (2018).

35. Isaac, R. S. et al. Nucleosome breathing and remodeling constrain CRISPR-Cas9 function. Elife 5, e13450 (2016).

36. Vigouroux, A., Oldewurtel, E., Cui, L., Bikard, D. & van Teeffelen, S. Tuning dCas9’s ability to block transcription enables robust, noiseless knockdown of bacterial genes. Mol. Syst. Biol. 14, e7899 (2018).

37. Tsai, S. Q. et al. CIRCLE-seq: a highly sensitive in vitro screen for genome-wide CRISPR-Cas9 nuclease off-targets. Nat. Meth. 14, 607–614 (2017).

38. Paquet, D. et al. Efficient introduction of specific homozygous and heterozygous mutations using CRISPR/Cas9. Nature 533, 125–129 (2016).

39. Richardson, C. D., Ray, G. J., DeWitt, M. A., Curie, G. L. & Corn, J. E. Enhancing homology-directed genome editing by catalytically active and inactive CRISPR-Cas9 using asymmetric donor DNA. Nat. Biotechnol. 34, 339–344 (2016).

40. Hu, J. H. et al. Evolved Cas9 variants with broad PAM compatibility and high DNA specificity. Nature 556, 57–63 (2018).

41. Kim, S., Kim, D., Cho, S. W., Kim, J. & Kim, J.-S. Highly efficient RNA-guided genome editing in human cells via delivery of purified Cas9 ribonucleoproteins. Genome Research 24, 1012–1019 (2014).

42. Kabadi, A. M., Ousterout, D. G., Hilton, I. B. & Gersbach, C. A. Multiplex CRISPR/Cas9-based genome engineering from a single lentiviral vector. Nucleic Acids Res. 42, e147–e147 (2014).

43. Gu, B. et al. Transcription-coupled changes in nuclear mobility of mammalian cis-regulatory elements. Science 359, 1050–1055 (2018).

44. Qi, L. S. et al. Repurposing CRISPR as an RNA-Guided Platform for Sequence-Specific Control of Gene Expression. Cell 152, 1173–1183 (2013).

45. Gilbert, L. A. et al. CRISPR-Mediated Modular RNA-Guided Regulation of Transcription in Eukaryotes. Cell 154, 442–451 (2013).

46. Kulcsár, P. I. et al. Crossing enhanced and high fidelity SpCas9 nucleases to optimize specificity and cleavage. Genome Biol 18, 347 (2017).

47. Casini, A. et al. A highly specific SpCas9 variant is identified by in vivo screening in yeast. Nat. Biotechnol. 36, 265–271 (2018).

48. McKenna, A. et al. Whole-organism lineage tracing by combinatorial and cumulative genome editing. Science 353, aaf7907 (2016).

49. Anders, C. & Jinek, M. In vitro enzymology of Cas9. Meth. Enzymol. 546, 1–20 (2014).

