## Supplementary Information for "Suppression of unwanted CRISPR/Cas9 editing by co-administration of catalytically inactivating truncated guide RNAs"

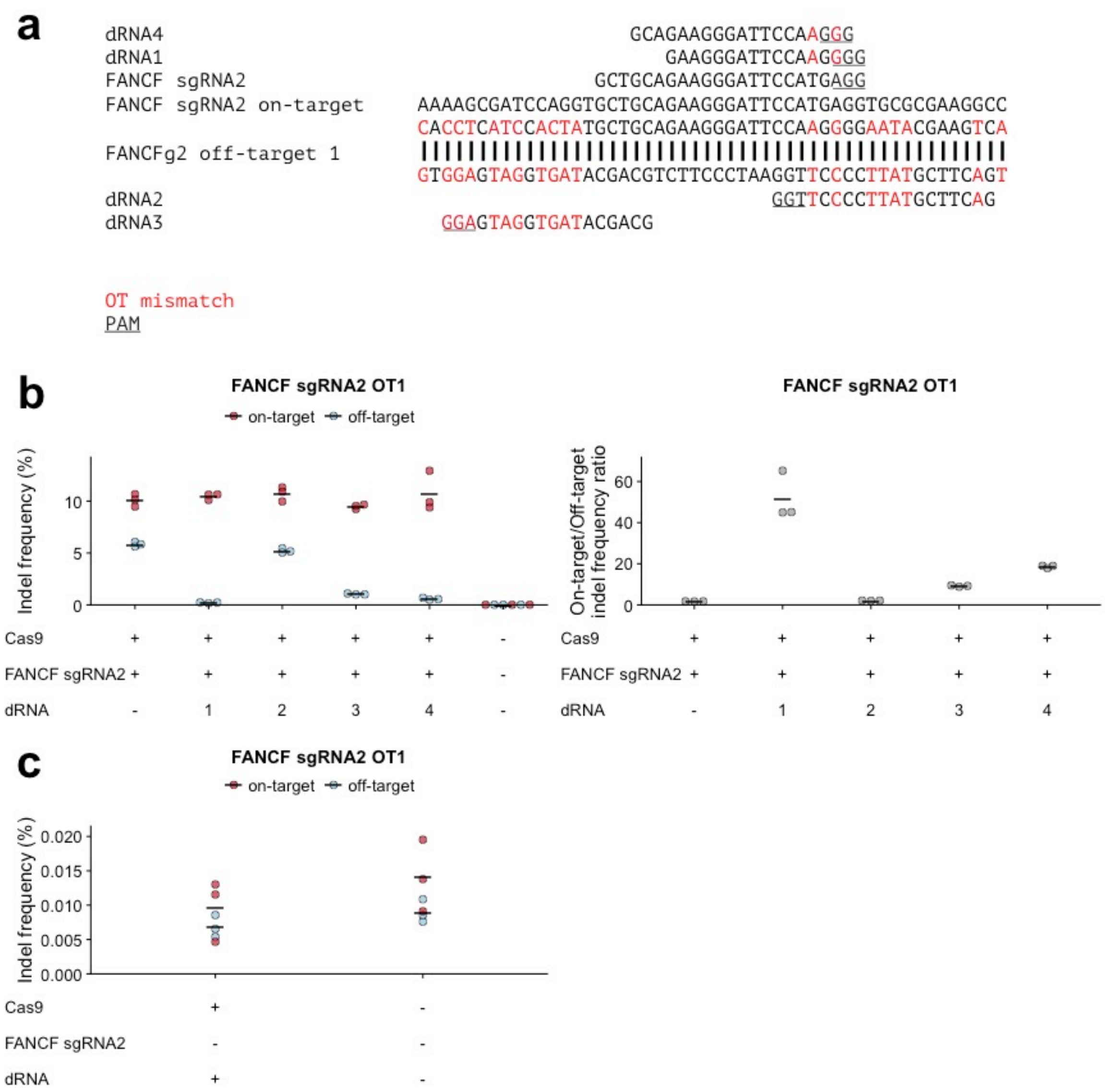


**Figure S1: FANCF dRNA1 does not promote Cas9-mediated editing. (a)** Sequence alignment of FANCF sgRNA2, its on-target site, the most prominent off-target, off-target site 1 (OT1), and multiple dRNAs complementary to OT1. **(b)** Indel frequencies and specificity ratios (on-target/off-target indel frequency ratios) at the FANCF sgRNA2 on-target site and OT1 24 hours after transfection with Cas9, sgRNA, and various dRNAs. For conditions without dRNA, an equivalent amount of pMAX-GFP was substituted. **(c)** Indel frequencies at the FANCF sgRNA2 on-target and OT1 sites 24 hours after transfection with Cas9 and dRNA1 but no sgRNA. The predicted cut sites of dRNA1 are the same as FANCF sgRNA2. Indel frequencies for untransfected cells are shown as a control. Numbers denote dRNA identity, see **Supplementary Data Set 1**. Solid lines denote the mean of n = 3 biological replicates.

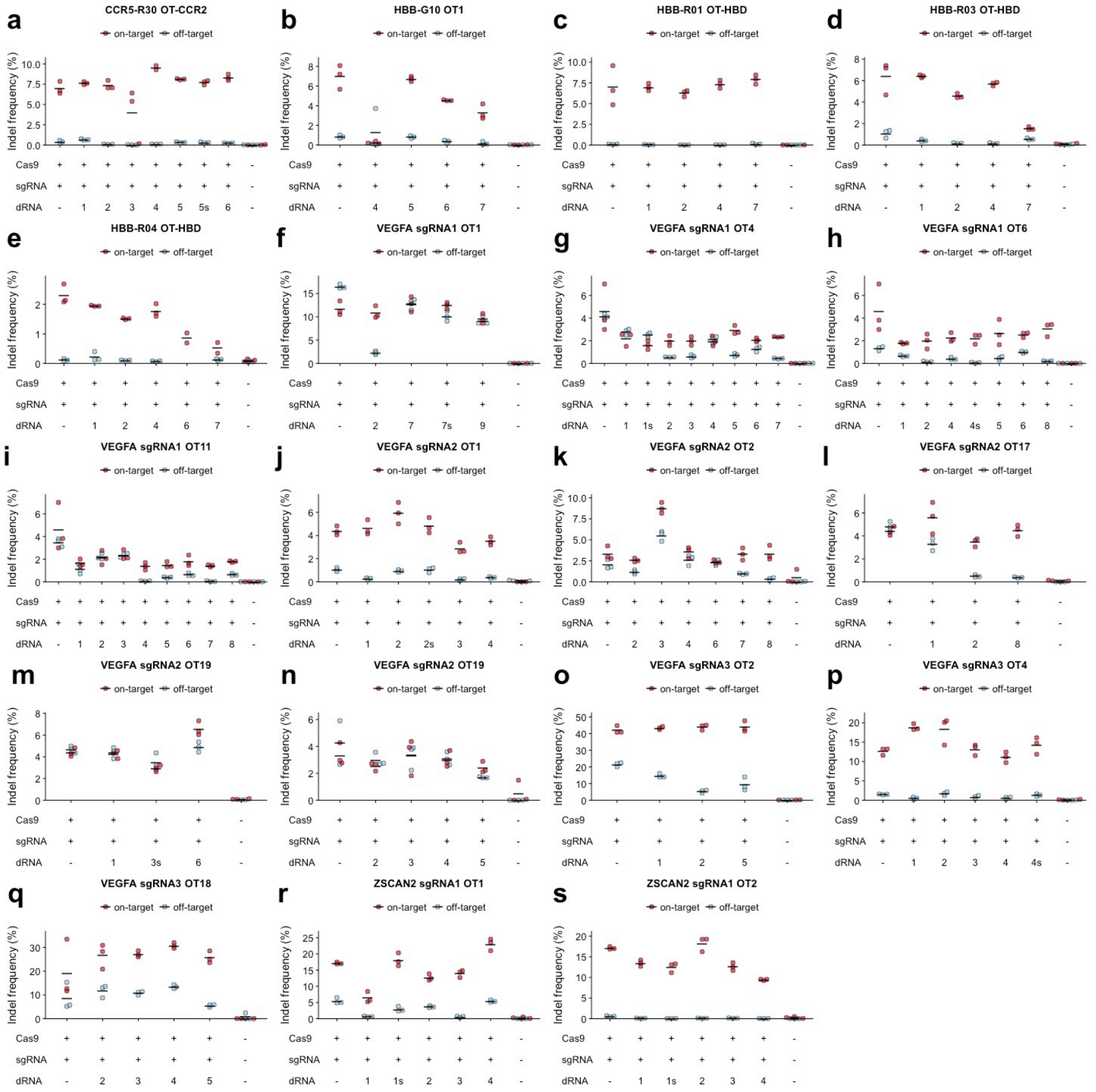

**Figure S2: dRNAs screened to increase the specificity ratios of 18 additional on-target/off-target pairs.** On and off-target indel frequencies 24 hours after transfection with Cas9, sgRNA, and off-target specific dRNAs in HEK293T cells **(a)** CCR5-R30 OT (CCR2). **(b)** HBB-G10 OT1. 4 additional dRNAs were screened, which are not shown here. **(c)** HBB-R01 OT (HBD). **(d)** HBB-R03 OT (HBD). **(e)** HBB-R04 OT (HBD). **(f)** VEGFA sgRNA1 OT1. **(g)** VEGFA sgRNA1 OT4. **(h)** VEGFA sgRNA1 OT6. **(i)** VEGFA sgRNA1 OT11. **(j)** VEGFA sgRNA2 OT1. **(k)** VEGFA sgRNA2 OT2. **(l)** VEGFA sgRNA2 OT17. **(m)** VEGFA sgRNA2 OT19 (dRNAs 1, 3s, and 6). **(n)** VEGFA sgRNA2 OT19 (dRNAs 2-5). **(o)** VEGFA sgRNA3 OT2. **(p)** VEGFA sgRNA3 OT4. **(q)** VEGFA sgRNA3 OT18. **(r)** ZSCAN2 sgRNA1 OT1. **(s)** ZSCAN2 sgRNA1 OT2. Indel frequencies for untransfected cells are shown as a control. Numbers denote dRNA identity, see **Supplementary Data Set 1**. Solid lines denote the mean of n = 3 biological replicates. OT = off-target.

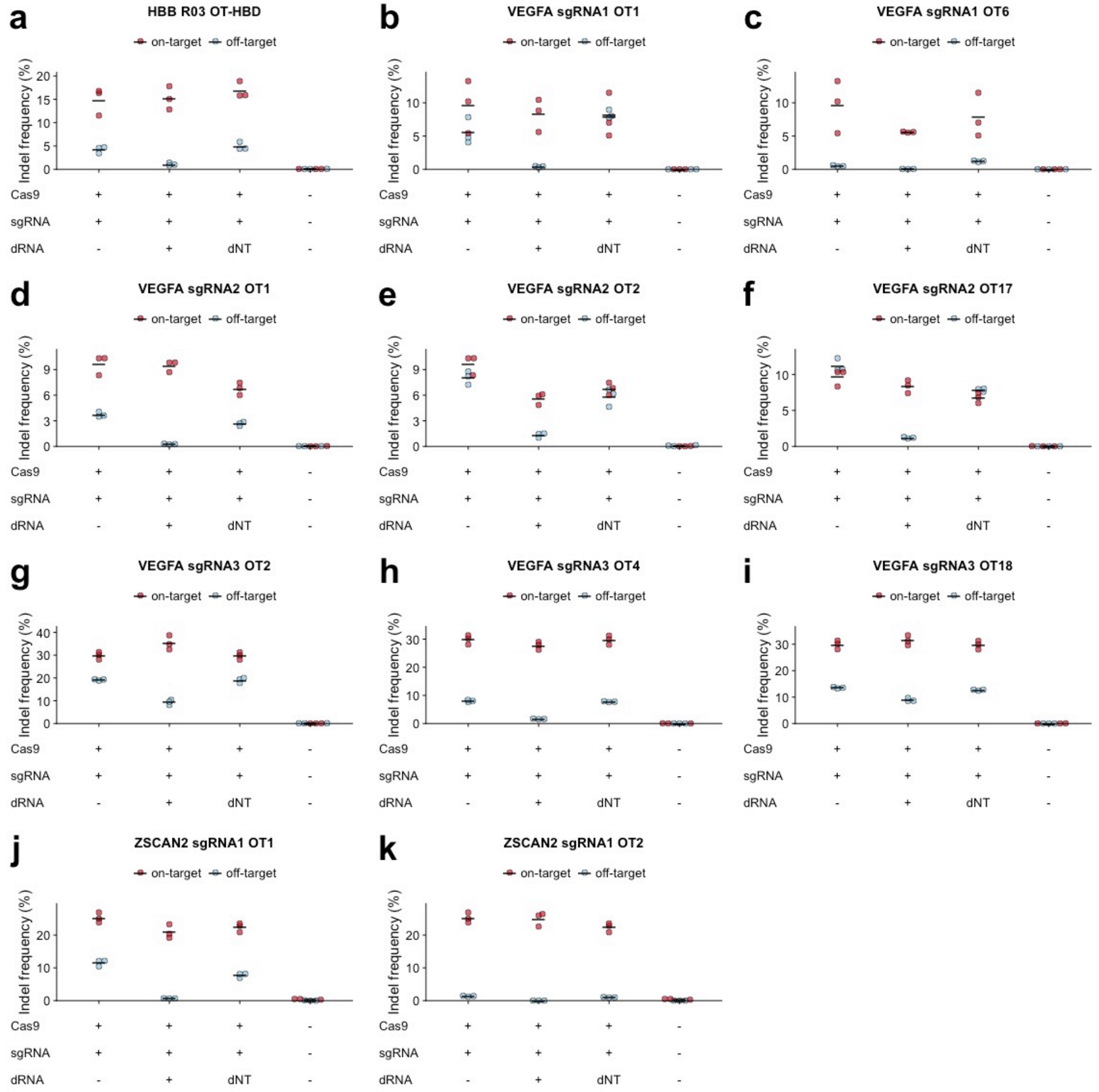


**Figure S3. Nontargeting dRNAs have minimal effects on on-target and off-target editing.** Comparison of most effective dRNA for 12 different off-target loci with a nontargeting dRNA (dNT). Indel frequency of on-target and off-target loci 24 hours after transfection with Cas9, sgRNA, ± dRNA or nontargeting dRNA in HEK293T cells. Indel frequencies for untransfected cells are shown as a control. **(a)** HBB R03 OT-HBD. **(b)** VEGFA sgRNA1 OT1. **(c)** VEGFA sgRNA1 OT6. **(d)** VEGFA sgRNA2 OT1. **(e)** VEGFA sgRNA2 OT2. **(f)** VEGFA sgRNA2 OT17. **(g)** VEGFA sgRNA3 OT2. **(h)** VEGFA sgRNA3 OT4. **(i)** VEGFA sgRNA3 OT18. **(j)** ZSCAN2 sgRNA1 OT1. **(k)** ZSCAN2 sgRNA1 OT2. Numbers denote dRNA identity, see **Supplementary Data Set 1**. Solid lines denote the mean of n = 3 biological replicates. OT = off-target.

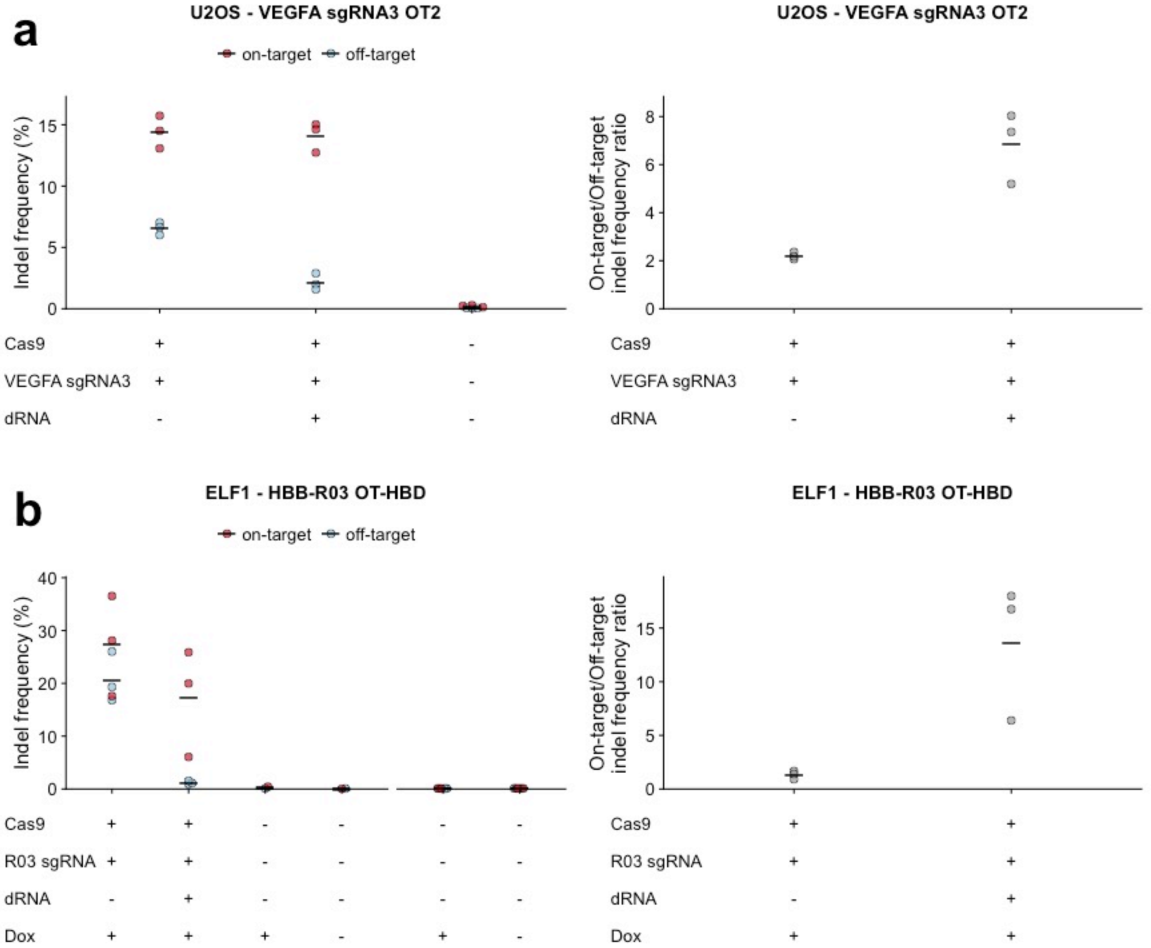


**Figure S4. dOTS is effective in multiple cell types.** On-target and off-target indel frequencies and specificity ratios 24 hours after transfection with Cas9, sgRNA, and off-target specific dRNA. **(a)** VEGFA sgRNA3 OT2 in U2OS cells. **(b)** HBB R03 OT (HBD) in Elf1 cells. Indel frequencies for untransfected cells are shown as a control. Control samples to the right of the x-axis break were performed separately. Solid lines denote the mean of n = 3 biological replicates. OT = off-target.

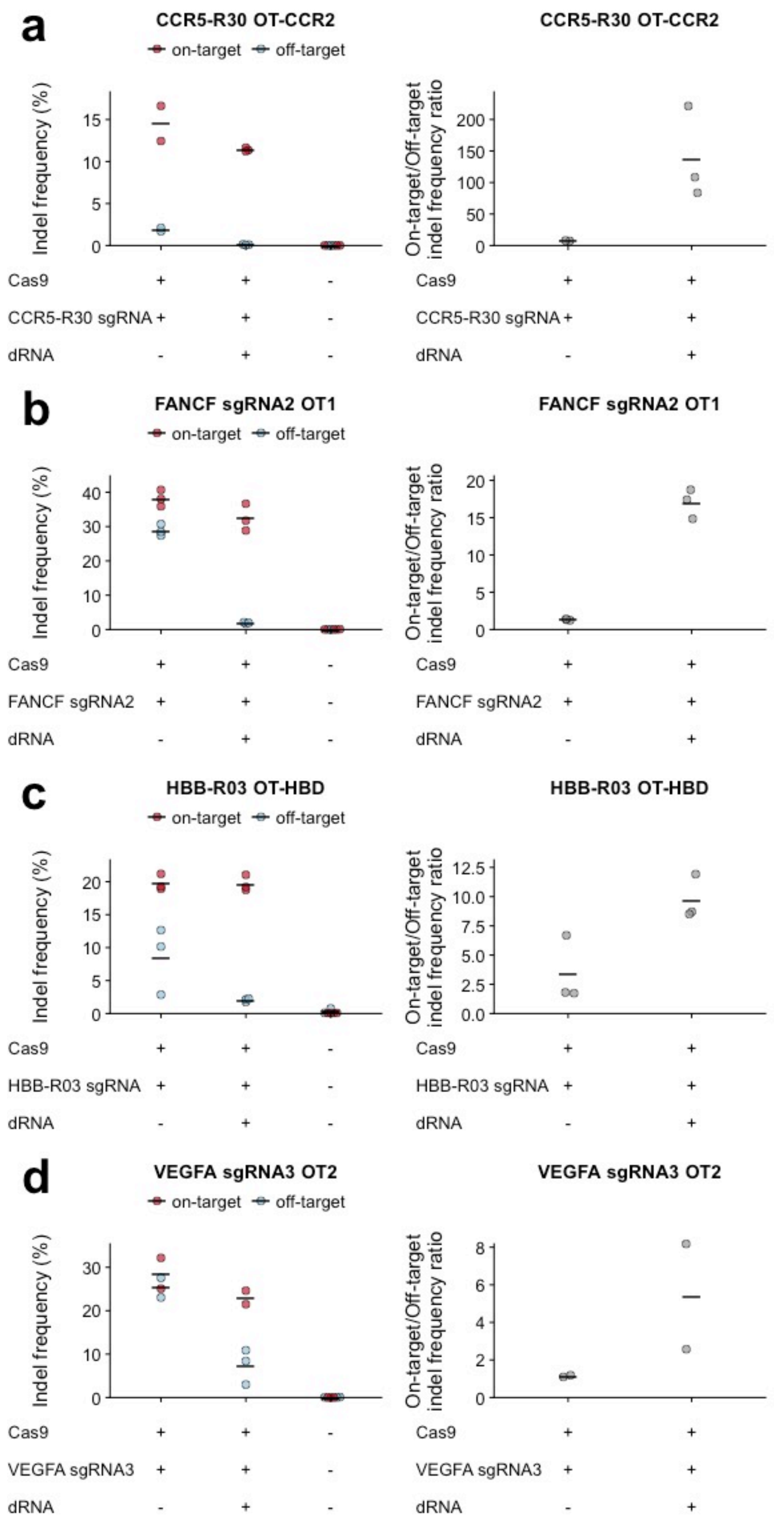


**Figure S5: dRNA-mediated off-target editing suppression is durable.** On-target and off-target indel frequencies and specificity ratios 72 hours after transfection with Cas9, sgRNA, and off-target specific dRNAs in HEK293T cells **(a)** CCR5-R30 OT (CCR2). **(b)** FANCF sgRNA2 OT1. **(c)** HBB-R03 OT (HBD). **(d)** VEGFA sgRNA3 OT2. Indel frequencies for untransfected cells are shown as a control. Numbers denote dRNA identity, see **Supplementary Data Set 1**. Solid lines denote the mean of n = 3 biological replicates, except CCR5-R30 and VEGFA sgRNA3 without dRNA where n = 2. 24 hour comparison shown in **Figure S2**.

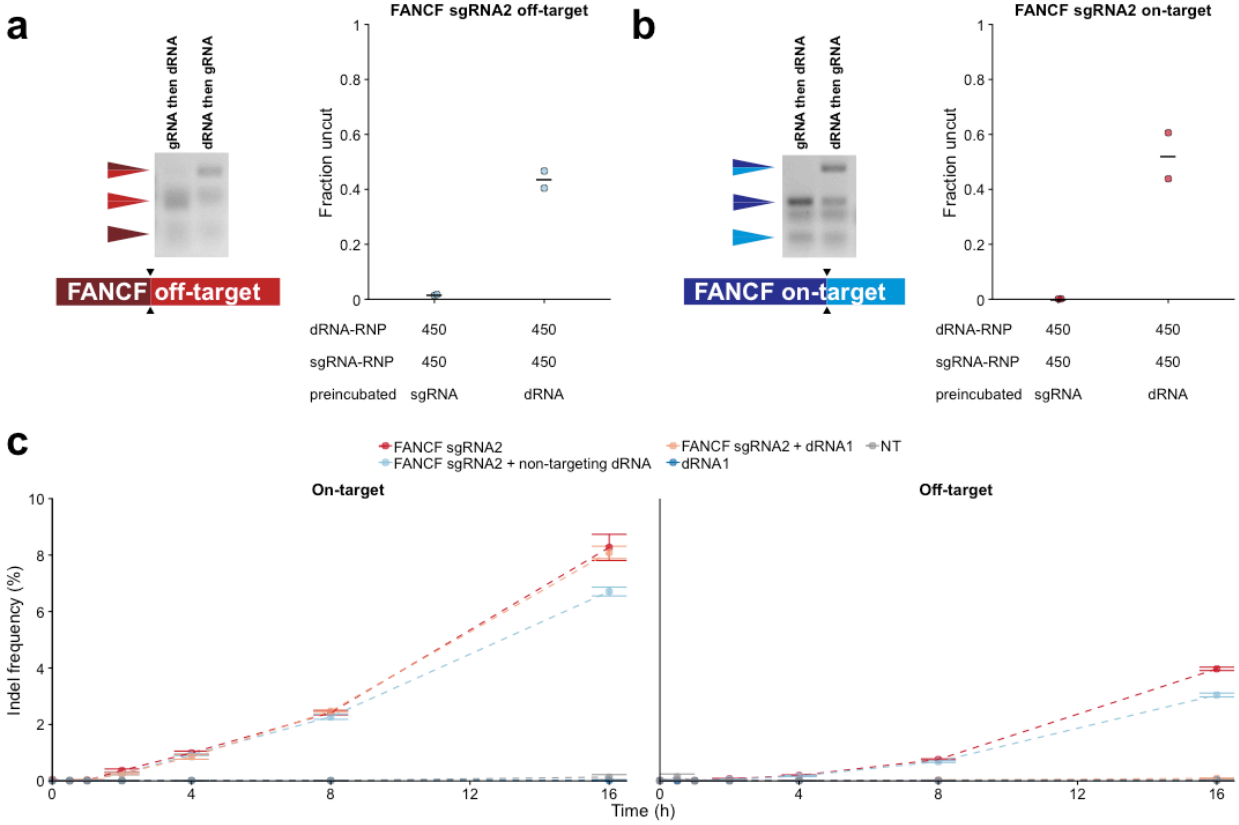


**Figure S6. dRNAs and sgRNAs compete for target site occupancy. (a, b)** Representative gels of *in vitro* Cas9 FANCF sgRNA2 RNP cleavage of linear PCR products containing either **(a)** the FANCF sgRNA2 off-target site (OT1) or **(b)** the FANCF sgRNA2 on-target site. PCR products were either preincubated with the sgRNA-RNP complex for 10 minutes prior to addition of the dRNA-RNP complex (sgRNA then dRNA) or were preincubated with the dRNA-RNP complex for 10 minutes prior to addition of the sgRNA-RNP complex (dRNA then sgRNA). ImageJ was used to quantify the intensity of the uncut and all cut bands in each lane. Fraction uncut was determined by dividing uncut intensity by sum of all band intensities in each lane. Solid lines denote the mean of n = 2 biological replicates. **(c)** Editing of FANCF on-target and off-target (OT1) sites in HEK293T cells using a chemically inducible Cas9 (ciCas9) from 0 to 16 hours after activation of ciCas9 with A115. A non-targeting dRNA was included as a control. NT = non-transfected control. Points depict the mean of n = 3 biological replicates, error bars depict the standard error of the mean.

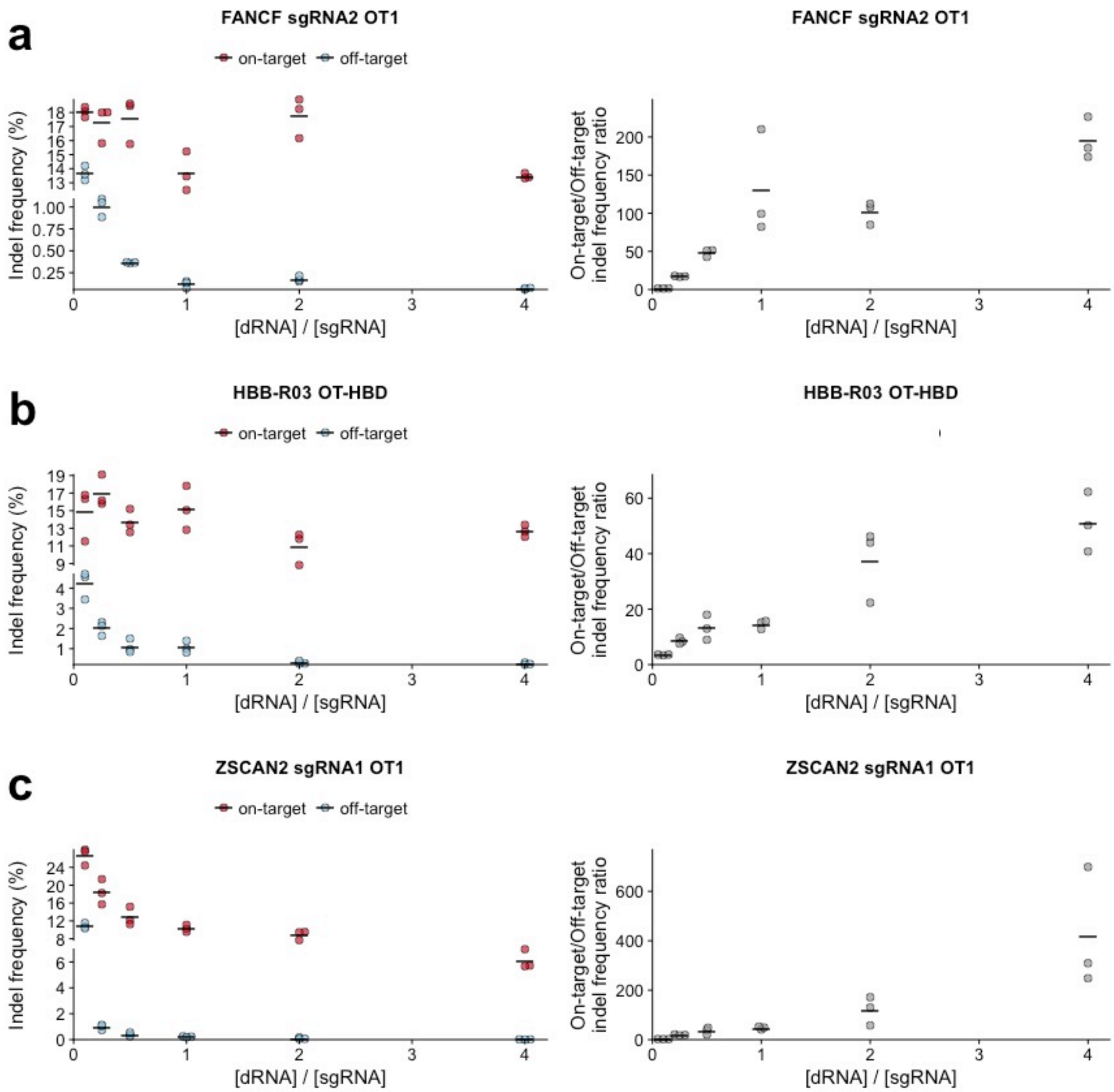


**Figure S7. Titration of dRNAs further reduces unwanted off-target editing at additional sites.** Target and off-target indel frequencies and specificity ratios 24 hours after transfection of various dRNA/sgRNA plasmid ratios for **(a)** FANCF sgRNA 2 and dRNA1; **(b)** HBB R03 and dRNA4; **(c)** ZSCAN2 sgRNA1 and dRNA3. Solid lines denote the mean of n = 3 biological replicates. OT = off-target.

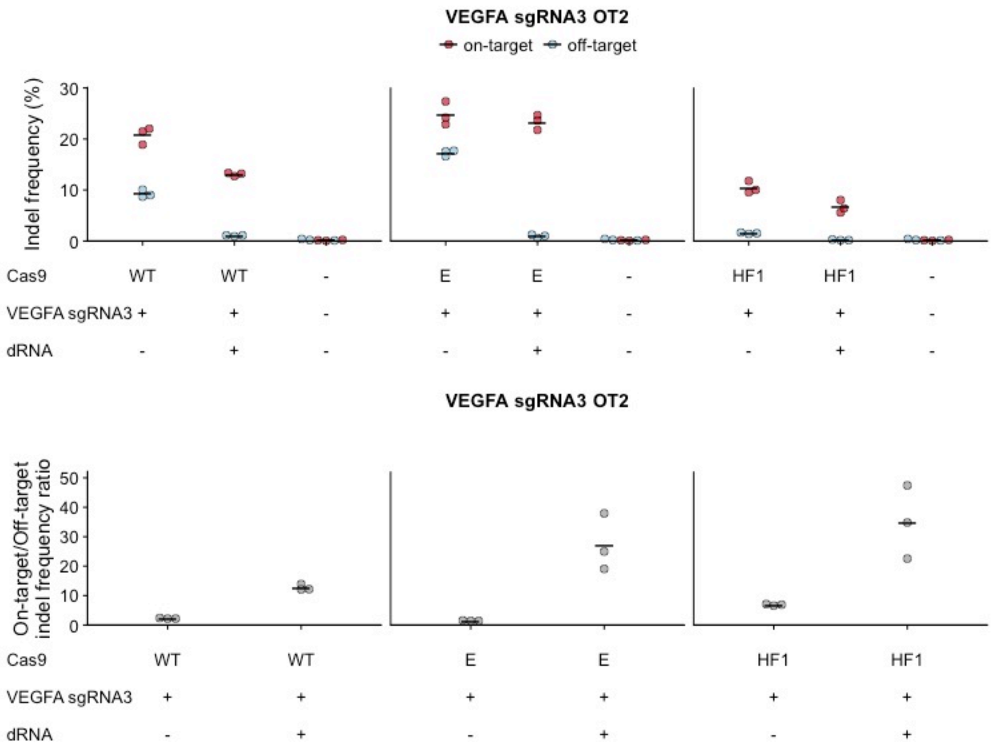


**Figure S8. dOTS can suppress refractory off-target editing of high-specificity Cas9 variants.** On-target and off-target indel frequencies and specificity ratios 24 hours after transfection of plasmids encoding VEGFA sgRNA3, dRNA and either wildtype Cas9 (WT), eSpCas9 (E), or SpCas9-HF1 (HF1). Indel frequencies for untransfected cells are shown as a control. Numbers denote dRNA identity, see **Supplementary Data Set 1**. Solid lines denote the mean of n = 3 biological replicates. OT = off-target.

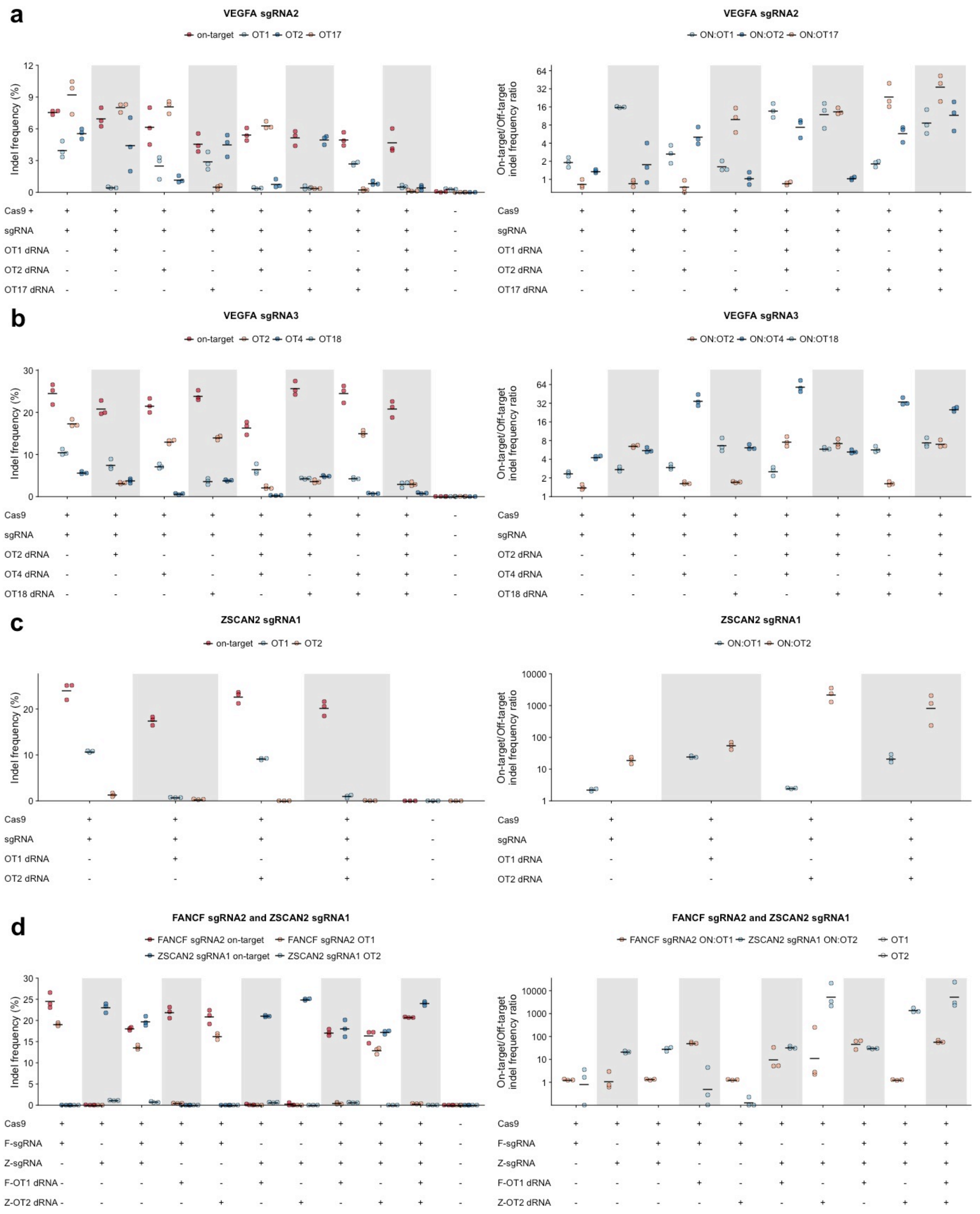


**Figure S9. dRNA can be combined to suppress unwanted off-target editing at a variety of sites.** Target and off-target indel frequencies and specificity ratios 24 hours after transfection with plasmids encoding Cas9 and various combinations of sgRNAs and dRNAs at **(a)** VEGFA sgRNA2

OT1, OT2, and OT17; **(b)** VEGFA sgRNA3 OT2, OT4, and OT18; **(c)** ZSCAN2 sgRNA1 OT1 and OT2; **(d)** FANCF sgRNA2 OT1 and ZSCAN2 sgRNA1 OT2. Indel frequencies for untransfected cells are shown as a control. Solid lines denote the mean of n = 3 biological replicates. OT = off-target.

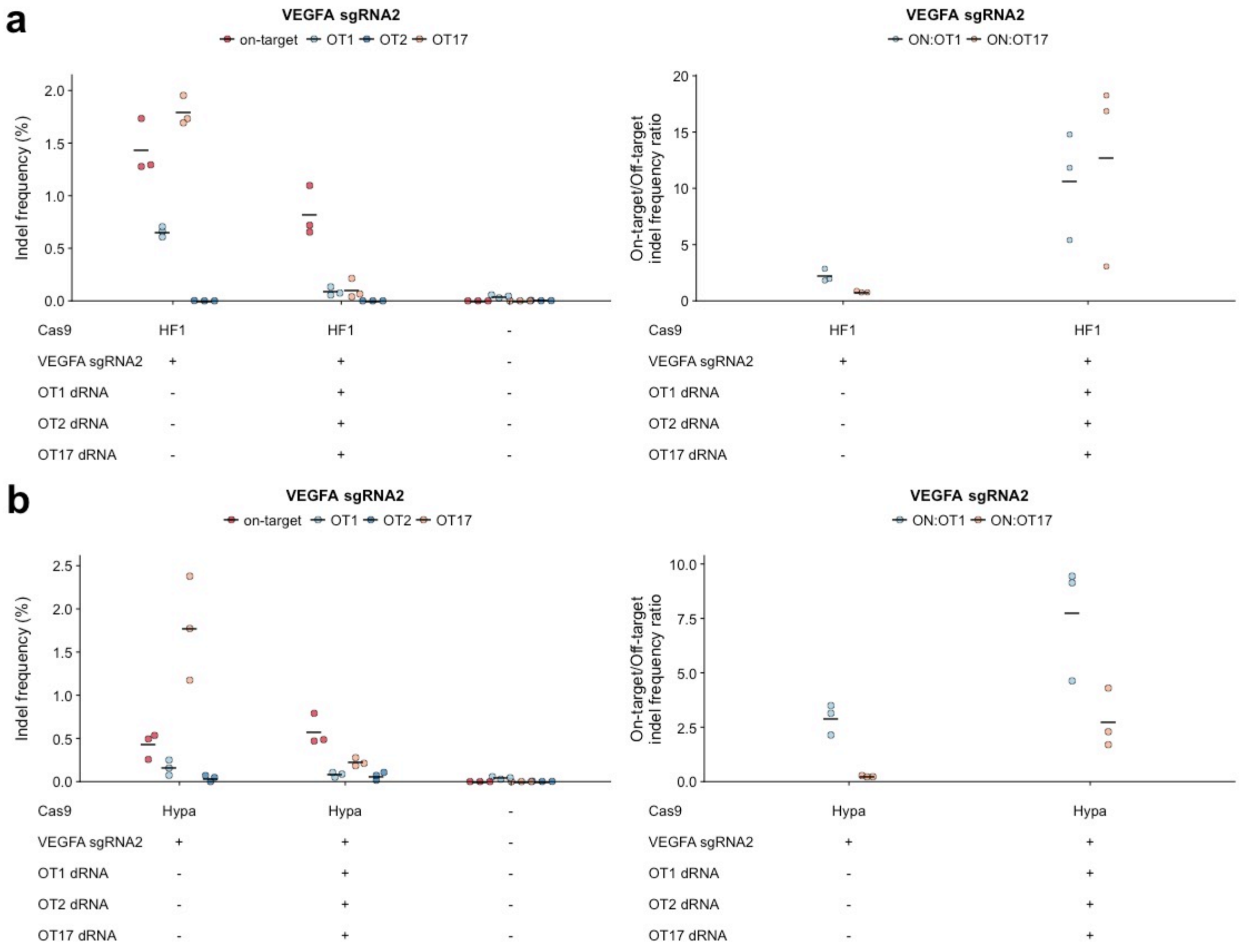


**Figure S10. Multiple dRNAs can be combined to reduce unwanted editing at multiple refractory off-target sites of high-specificity Cas9 variants.** Target and off-target indel frequencies and specificity ratios 24 hours after transfection with plasmids encoding VEGFA sgRNA2, a combination of three dRNAs and either **(a)** SpCas9-HF1 (HF1) or **(b)** HypaCas9 (Hypa). Despite being reported previously[21], indels were not observed at OT2, so specificity ratios were not plotted. Indel frequencies for untransfected cells are shown as a control. Solid lines denote the mean of n = 3 biological replicates. OT = off-target.

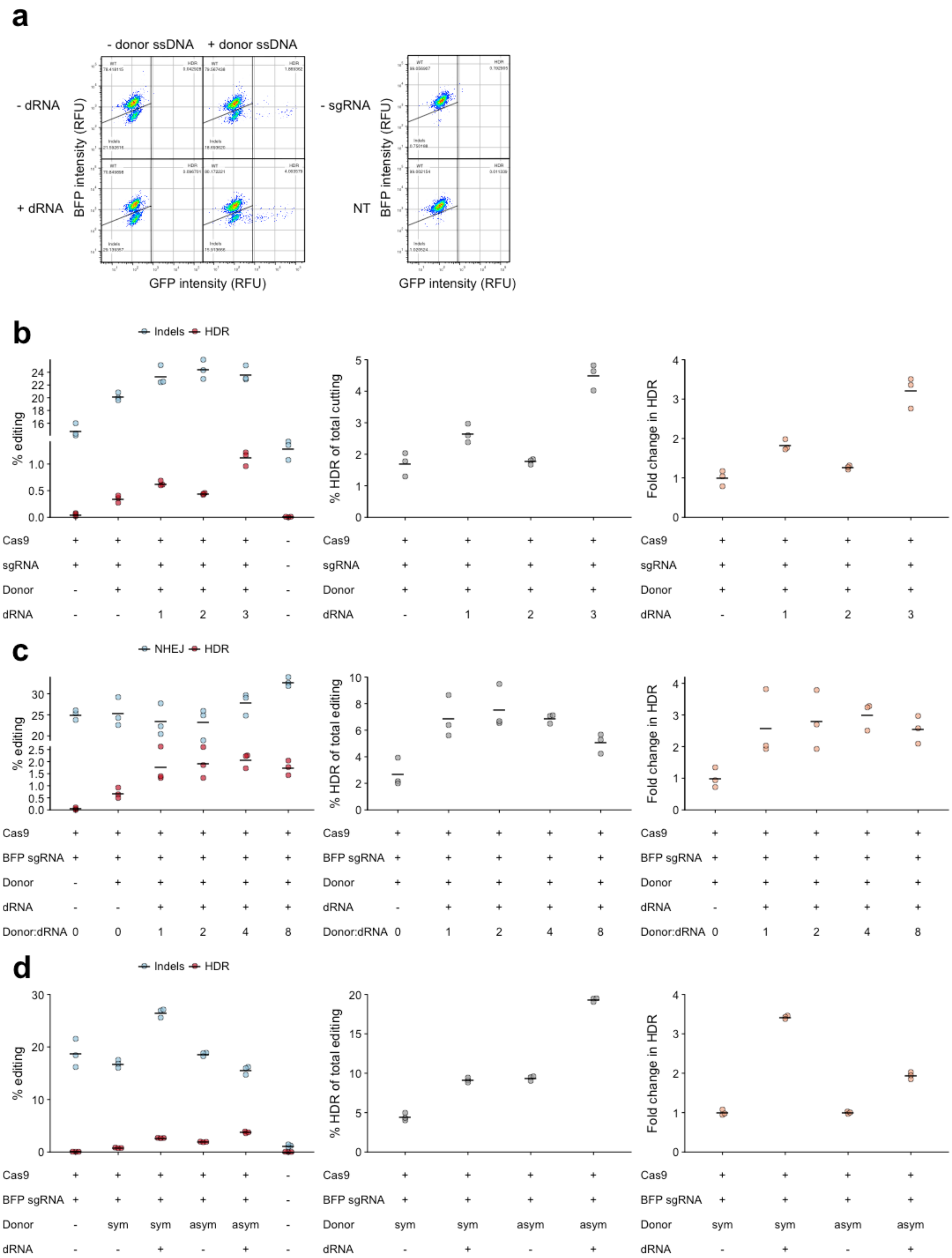
a
- donor ssDNA
+ donor ssDNA
- dRNA
+ dRNA
BFP intensity (RFU)
GFP intensity (RFU)
- sgRNA
NT
b
Indels
HDR
% editing
% HDR of total cutting
Fold change in HDR
Cas9
sgRNA
Donor
dRNA
c
NHEJ
HDR
% editing
% HDR of total editing
Fold change in HDR
Cas9
BFP sgRNA
Donor
dRNA
Donor:dRNA
d
Indels
HDR
% editing
% HDR of total editing
Fold change in HDR
Cas9
BFP sgRNA
Donor
dRNA
sym
asym

**Figure S11. Screening guides, donors, and dRNAs for scarless HDR in a fluorescent reporter system. (a)** Representative flow cytometry plots illustrating alteration of a synthetic BFP locus to GFP after Cas9•sgRNA editing. **(b)** Screening of three dRNAs for indels or HDR events, percent HDR of total Cas9 editing observed, and fold change in HDR observed. **(c)** Screening of various ratios of dRNA3 to sgRNA for NHEJ or HDR events, percent HDR of total Cas9 editing observed, and fold change in HDR observed. **(d)** Comparison of symmetric donor and asymmetric donor for NHEJ or HDR events, percent HDR of total Cas9 editing observed, and fold change in HDR observed. HDR donors do not contain blocking mutations. Indel frequencies for untransfected cells are shown as a control. Numbers denote sgRNA and dRNA identities, see **Supplementary Data Set 1**. Solid lines denote the mean of n = 3 biological replicates.

### Supplmentary Tables

| Site | Best dRNA | n | Normalized specificity ratio (mean) | On-target ratio (mean) | Normalized specificity ratio (s.e.m) | On-target ratio (s.e.m) |
|---|---|---|---|---|---|---|
| **ZSCAN2 sgRNA1 OT2** | 1s | 3 | 37.93 | 0.73 | 7.07 | 0.04 |
| **FANCF sgRNA2 OT1** | 1 | 3 | 29.96 | 1.04 | 3.42 | 0.04 |
| **VEGFA sgRNA2 OT17** | 8 | 3 | 13.11 | 1.02 | 1.57 | 0.08 |
| **CCR5-R30 OT-CCR2** | 3 | 3 | 11.34 | 0.57 | 6.81 | 0.28 |
| **ZSCAN2 sgRNA1 OT1** | 3 | 3 | 8.95 | 0.82 | 2.07 | 0.04 |
| **VEGFA sgRNA2 OT2** | 8 | 3 | 7.50 | 1.00 | 2.60 | 0.21 |
| **HBB-R01 OT-HBD** | 2 | 3 | 7.48 | 0.89 | 1.74 | 0.18 |
| **VEGFA sgRNA3 OT4** | 1 | 3 | 6.75 | 1.48 | 1.68 | 0.07 |
| **VEGFA sgRNA1 OT1** | 2 | 3 | 6.72 | 0.93 | 0.97 | 0.10 |
| **HBB-R03 OT-HBD** | 4 | 3 | 6.55 | 0.89 | 2.01 | 0.12 |
| **VEGFA sgRNA1 OT4** | 8 | 3 | 4.99 | 0.51 | 1.37 | 0.14 |
| **VEGFA sgRNA1 OT6** | 7 | 3 | 4.57 | 0.66 | 1.49 | 0.19 |
| **VEGFA sgRNA2 OT1** | 1 | 3 | 4.32 | 1.05 | 0.92 | 0.10 |
| **VEGFA sgRNA3 OT2** | 2 | 3 | 4.26 | 1.04 | 0.43 | 0.04 |
| **VEGFA sgRNA3 OT18** | 5 | 3 | 2.13 | 1.33 | 1.13 | 0.50 |
| **VEGFA sgRNA1 OT11** | 7 | 3 | 40.60 | 0.31 | 16.94 | 0.08 |
| **HBB-G10 OT1** | 7 | 3 | 3.74 | 0.47 | 1.49 | 0.08 |
| **VEGFA sgRNA2 OT19** | 5 | 3 | 1.55 | 0.72 | 0.55 | 0.13 |
| **HBB-R04 OT-HBD** | 4 | 3 | 1.16 | 0.77 | 0.29 | 0.09 |

**Supplementary Table 1 | dRNAs designed for a variety of sites increase specificity ratio with minimal effects on on-target editing.** Normalized specificity ratios, computed as the specificity ratio in the presence of the best dRNA at a site divided by the specificity ratio in the absence of the dRNA, and on-target ratios, computed as the ratio of on-target editing in the presence of the best dRNA at a site divided by the on-target editing in the absence of the dRNA, for the best dRNA for 19 sgRNA/off-target pairs. n = 3 biological replicates, error measured as the standard error of the mean (s.e.m.).

s

| **Site** | **Δ (On)** | **p (On)** | **$p_{adj}$ (On)** | **Δ (OT)** | **p (OT)** | **$p_{adj}$ (OT)** |
|---|---|---|---|---|---|---|
| **FANCF sgRNA2 OT1** | -0.004 | 0.835 | 1 | -0.002 | 0.910 | 1 |
| **HBB R03 OT-HBD** | -0.014 | 0.845 | 1 | -0.009 | 0.761 | 1 |
| **VEGFA sgRNA1 OT1** | 6.07E-04 | 0.403 | 1 | 6.61E-04 | 0.094 | 1 |
| **VEGFA sgRNA1 OT6** | -1.95E-04 | 0.524 | 1 | 0.025 | 0.209 | 1 |
| **VEGFA sgRNA2 OT1** | -0.045 | 0.912 | 1 | 0.083 | 0.124 | 1 |
| **VEGFA sgRNA2 OT2** | -0.018 | 0.750 | 1 | -9.37E-04 | 0.683 | 1 |
| **VEGFA sgRNA2 OT17** | 0.015 | 0.306 | 1 | 0.008 | 0.218 | 1 |
| **VEGFA sgRNA3 OT4** | -0.007 | 0.907 | 1 | 0 | 1 | 1 |
| **VEGFA sgRNA3 OT18** | -3.27E-04 | 0.513 | 1 | 0.002 | 0.319 | 1 |
| **ZSCAN2 sgRNA1 OT1** | -3.87E-04 | 0.789 | 1 | 0 | 1 | 1 |
| **ZSCAN2 sgRNA1 OT2** | 3.15E-05 | 0.479 | 1 | 0.001 | 0.092 | 1 |
| **VEGFA sgRNA3 OT2** | 0.040 | 0.406 | 1 | 0.080 | 0.050 | 1 |

**Supplementary Table 2 | dRNAs alone do not promote editing at sgRNA target sites..** Difference between indel frequencies at on- and off-target (OT) sites for the best dRNA at 12 different on/off-target pairs (Δ). p: p-value, based on two-sided Student's t-test. $p_{adj:}$ Bonferroni-adjusted p-value. n = 3 biological replicates at on- and off-target sites, except for VEGFA sgRNA3 OT2 (n = 9) and VEGFA sgRNA3 OT18 (n = 3 at on-target, n = 2 at off-target due to failed sequencing reactions).

| Site | Δ ($On_{pred}$) | p ($On_{pred}$) | $p_{adj}$ ($On_{pred}$) | Δ ($OT_{pred}$) | p ($OT_{pred}$) | $p_{adj}$ ($OT_{pred}$) |
|---|---|---|---|---|---|---|
| **FANCF sgRNA2 OT1** | -0.004 | 0.835 | 1 | -0.002 | 0.910 | 1 |
| **HBB R03 OT-HBD** | -0.056 | 0.699 | 1 | -0.006 | 0.828 | 1 |
| **VEGFA sgRNA1 OT1** | -0.004 | 0.730 | 1 | 0.001 | 0.312 | 1 |
| **VEGFA sgRNA1 OT6** | 7.78E-04 | 0.278 | 1 | 0.027 | 0.204 | 1 |
| **VEGFA sgRNA2 OT1** | 0.316 | 0.220 | 1 | 0.083 | 0.124 | 1 |
| **VEGFA sgRNA2 OT2** | 0.028 | 0.253 | 1 | 0.001 | 0.302 | 1 |
| **VEGFA sgRNA2 OT17** | 0.092 | 0.115 | 1 | 0.074 | 0.260 | 1 |
| **VEGFA sgRNA3 OT4** | -0.015 | 0.908 | 1 | 0 | 1 | 1 |
| **VEGFA sgRNA3 OT18** | 6.55E-05 | 0.471 | 1 | -0.013 | 0.622 | 1 |
| **ZSCAN2 sgRNA1 OT1** | 0.002 | 0.089 | 1 | -7.37E-04 | 0.539 | 1 |
| **ZSCAN2 sgRNA1 OT2** | 3.15E-05 | 0.479 | 1 | 0.001 | 0.092 | 1 |
| **VEGFA sgRNA3 OT2** | 0.061 | 0.094 | 1 | 0.073 | 0.071 | 1 |

**Supplementary Table 3 | dRNAs alone do not promote editing at predicted dRNA target sites.** Difference between indel frequencies at on- and off-target (OT) sites for the best dRNA at 12 different on/off-target pairs (Δ). Predicted indel locations (pred) are the location of expected indels if the dRNA were a full length sgRNA. p: p-value, based on two-sided Student's t-test. $p_{adj:}$ Bonferroni-adjusted p-value. n = 3 biological replicates at on- and off-target sites, except for VEGFA sgRNA3 OT2 (n = 9) VEGFA sgRNA2 OT17 (n = 3 at on-target, n = 2 at off-target due to failed sequencing reactions), and VEGFA sgRNA3 OT18 (n = 3 at on-target, n = 2 at off-target due to failed sequencing reactions).
